# Anchors for Homology-Based Scaffolding

**DOI:** 10.1101/2025.04.28.650980

**Authors:** Karl K. Käther, Thomas Gatter, Steffen Lemke, Peter F. Stadler

## Abstract

Homology-based scaffolding orders contigs based on conserved collinearity of homologous sequences across related species. Existing methods often rely on costly whole-genome alignments or show limited robustness when integrating multiple references. Here, we introduce an anchor-based scaffolding framework that adapts synteny anchors to efficiently infer contig order and orientation relative to one or more reference genomes. Our approach leverages precomputed, sufficiently unique anchors and their respective high-confidence homology matches in a greedy approach, combining single-reference to multi-reference scaffolds using a maximum matching. Across simulated and real datasets, anchor-based scaffolding achieves accuracy comparable to state-of-the-art methods. Notably, the approach shows particular strengths in multi-reference settings. These results demonstrate that synteny-anchor-based scaffolding provides an additional tool for homology-based scaffolding with robust accuracy and superior performance in multi-reference scenarios.

## 1. Introduction

High-throughput sequencing and assembly pipelines have been well established in the last decades, allowing researchers to create high quality contigs for genomes of varied species. Yet, scaffolding, the task of arranging them in correct chromosomal order and orientation, remains a challenging and contested area of research. Scaffolding typically requires additional data, because paired-end reads alone do not possess sufficient discriminatory power. While Hi-C approaches have become the de facto gold standard ^1,2^, homology-based scaffolding using closely related reference genomes offers an attractive alternative as high-quality reference assemblies become increasingly available. Several approaches have been developed for this task ^3,4,5,6^, including tools for multi-reference (multi-ref) scaffolding ^7,8^.

Current homology-based scaffolding approaches face several challenges. Computationally, methods relying on whole-genome alignments can be resource-intensive; for example, Progressive Cactus alignments required for `Ragout2` ^6^ are extremely computationally demanding for large eukaryotic genomes ^5^. Biologically, repetitive and duplicated genomic regions complicate scaffolding by introducing noisy signal, while larger-scale rearrangements between target and reference add further complexity. Additionally, errors from earlier assembly steps can also introduce conflicting information. Employing multiple references amplifies these contradictions, requires phylogenetic information to weigh references appropriately, and demands additional computational resources.

Synteny anchors - sufficiently unique sequences forming unambiguous 1-1 orthologous matches ^9,10^ - offer solutions to these limitations. By construction, anchors exclude duplicated and many otherwise rearranged regions, reducing alignment ambiguity which promises to enable development of efficient algorithms for contig ordering.

In this contribution we adapt our recently developed `AncST` synteny anchor framework ^10^, originally designed for synteny detection, for the purpose of contig scaffolding. Formally, scaffolding assigns each contig *i* a position *π*(*i*) and orientation *o*(*i*) ∈ {−1, +1} in the final assembly, seeking to maximize a score *S*(*π, o*) that rewards collinearity with the reference. This optimization problem is NP-hard ^11^, motivating heuristic approaches. Our method employs a greedy algorithm that orders contigs based on anchor positions along reference chromosomes, then applies a maximum weighted perfect matching approach ^8^ to reconcile single-reference (single-ref) scaffolds into multi-ref consensus scaffolds. This work is timely given massive genome sequencing efforts such as the Darwin Tree of Life,^a^ the European Reference Genome Atlas,^b^ and the Earth Biogenome Project.^c^

## 2. Methods

### 2.1 Anchors

In recent work, we investigated the notion of annotation-free anchors as a means to identify syntenic regions efficiently; see ^9,10^ for full detail. In brief, a subsequence *w* of a genome is called *d*_0_-unique if *d*(*w, x*) *> d*_0_ holds for every other subsequence *x* not overlapping *w*, and *d* denotes a metric pairwise sequence distance. Two *d*_0_-unique subsequences *w* and *w*^*′*^ in different genomes in *G* and *G*^*′*^ are guaranteed to form a unique match if *d*(*w, w*^*′*^) *< d*_0_*/*2. In practice, it suffices to use a cutoff of *d*_0_ minus some small tolerance. We call *d*_0_-unique sequences anchor candidates. Pairs of anchor candidates *w* and *w*^*′*^ satisfying 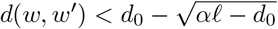 ^10^ form *anchors* between the genomes *G* and *G*^*′*^.

Anchors were used in ^10^ to construct fine-grained synteny maps between not-too-distant genomes. In order to determine anchor candidates efficiently, we use infrequent *k*-mers with a length close to log_4_ |*G*|, where |*G*| denotes the genome size, as an indicator for the uniqueness of *w*. Details of the implementation are described in ^10^.

### 2.2 Contig Sorting

The basic idea underlying our approach is rather straightforward: Since anchors between target contigs and reference chromosomes establish a robust 1-1 orthology relationship, they also anchor contigs at specific locations of the reference assembly. The same 1-1 relationship also justifies a simple greedy approach. Thus, we first determine, for each contig, the reference chromosome with the largest number of matched anchors. In the current implementation we do not attempt to identify genome rearrangements between reference and target explicitly. Instead, we directly construct homology-based scaffolds that (locally) match within chromosomes of the reference assembly. We will discuss this point in more detail in the Conclusions section.

After global construction of anchors, we consider each reference chromosome separately. Let ℒ be an initially empty list. We traverse the anchors between the reference chromosome and its assigned contigs in the order of the reference. For each contig, we keep count of the number of its anchors that have been visited so far. A contig *i* is appended to ℒ as soon as at least half of its anchors have been visited. The order of their insertion into ℒ therefore determines the positions *π*(*i*) of the contigs in the emerging scaffold. Their orientations *o*(*i*) are assigned based on the predominant strand direction of the anchor matches with the assigned reference sequence. The use of `AncST` anchors, i.e., nearly perfect 1-1 orthologs, makes the greedy approach a natural choice since overlapping matches of contigs to the references are expected to be absent in theory and extremely rare in practice. This simple greedy strategy for determining a contig order relative to a reference assembly can easily be extended to multiple reference assemblies by simply running anchor detection and greedy ordering independently for each reference assembly. Different reference assemblies will yield slightly different contig orders; it then remains to reconcile the conflicting order information.

### 2.3 Merging of Alternative Scaffolds

Scaffolds obtained from different reference assemblies can be reconciled using a simple graph-theoretic approach ^8^. To this end, we define a graph Γ = (*V, E, w*) whose vertex set are the end points of the target contigs, i.e., each target contig is represented by *two* vertices. An edge {*u, v*} is inserted if *u* and *v* are endpoints of two distinct contigs that are consecutive for one of the reference assemblies. Note that for a pair of consecutive contigs the choice of the adjacent endpoints determines the relative orientation of the two contigs. Each edge {*u, v*} has a weight *w* that in the simplest case counts the number of reference assemblies in which *u* and *v* are consecutive. More sophisticated weights *w* can be introduced that account e.g. for the quality of anchors for the contigs of which *u* and *v* are endpoints. Weights are aggregated if an edge appears for more than one reference assembly. For technical reasons, it is useful, moreover, to assign edges with weight 0 to all pairs of vertices that are neither consecutive w.r.t. at least one reference assembly nor form the endpoints of the same contig. This amounts to building a single component with circular connections.

In order to extract a consensus scaffold from the weighted graph Γ, we observe that in a scaffold each contig has at most one predecessor and one successor and thus each end-point of a scaffold can be incident to at most one edge. Hence scaffolds that only contain observed adjacencies correspond to matchings in Γ. A maximum weighted matching in Γ thus serves as a best estimate for a consensus scaffolding. We note that a matching will be perfect, i.e. includes all vertices, only if all contigs are placed into circular chromosomes. Otherwise, there will be contig ends that correspond to ends of a scaffold and thus are not incident to any edge. The maximum weighted matching problem can be solved in polynomial time using the *blossom* algorithm ^12^.

Adding missing edges to Γ with weight 0 transforms the problem into a maximum weight (or minimum cost) *perfect matching* problem, which can be solved by the highly optimized `Blossom V` implementation ^13^. This approach has been pioneered in ^8^; we refer to Figure 2 in this publication for a detailed visual description of the algorithm.

**Fig. 1.**
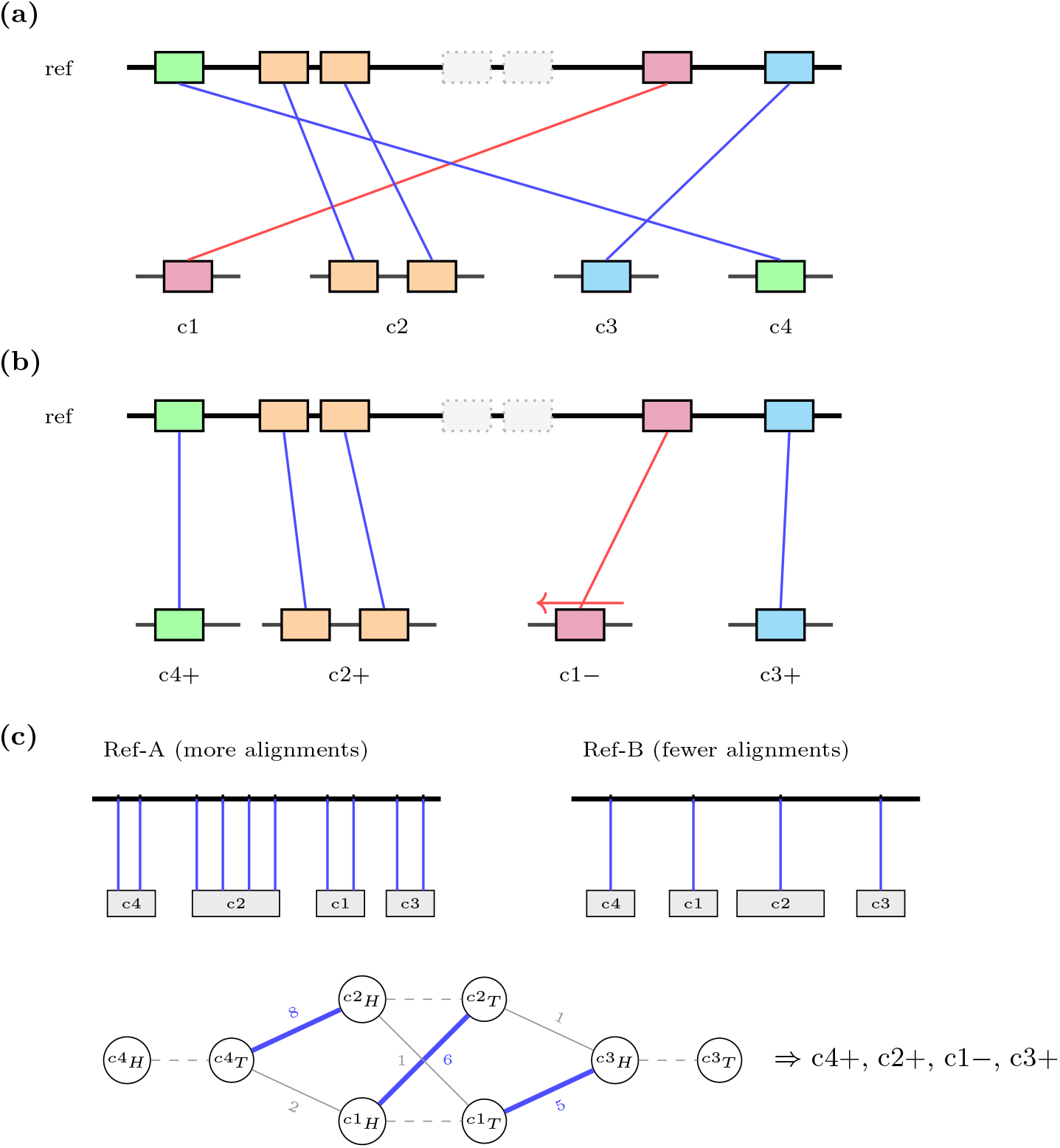
AncST pipeline. **(a)** *Anchor generation*. Each colored rectangle on the reference and on a contig is a *d*_0_-unique anchor candidate; matched candidates of identical color form an anchor pair (line between them). The two adjacent gray dotted rectangles on the reference are a tandem duplication not yielding an anchor because of low *d*_0_-uniqueness. The red link to *c*1 indicates its anchor matches on the opposite strand. Contigs (*c*1 – *c*4) are shown in their initial unordered state. **(b)** *Single-reference contig ordering*. Contigs are arranged greedily according to the position of their anchors along the reference; orientations *±* follow the dominant strand of the matches (red link plus inversion arrow above *c*1 records its reverse alignment). **(c)** *Multi-reference merging*. Two pairwise scaffolds, from Ref-A (denser anchor coverage) and Ref-B (sparser), disagree on the local order of *c*1 and *c*2. Each contig contributes a head (*H*) and tail (*T*) vertex with an implicit intra-contig connection (dashed). Pairwise adjacencies become weighted edges (weights illustrative; the *c*4_*T*_ –*c*2_*H*_ edge has weight 8, well above *c*4_*T*_ –*c*1_*H*_ at 2, because Ref-A supports it more strongly). Maximum-weight perfect matching (Blossom V) selects the high-weight Ref-A adjacencies (thick blue) over the conflicting Ref-B ones (thin gray), yielding the consensus order on the right.

**Fig. 2.**
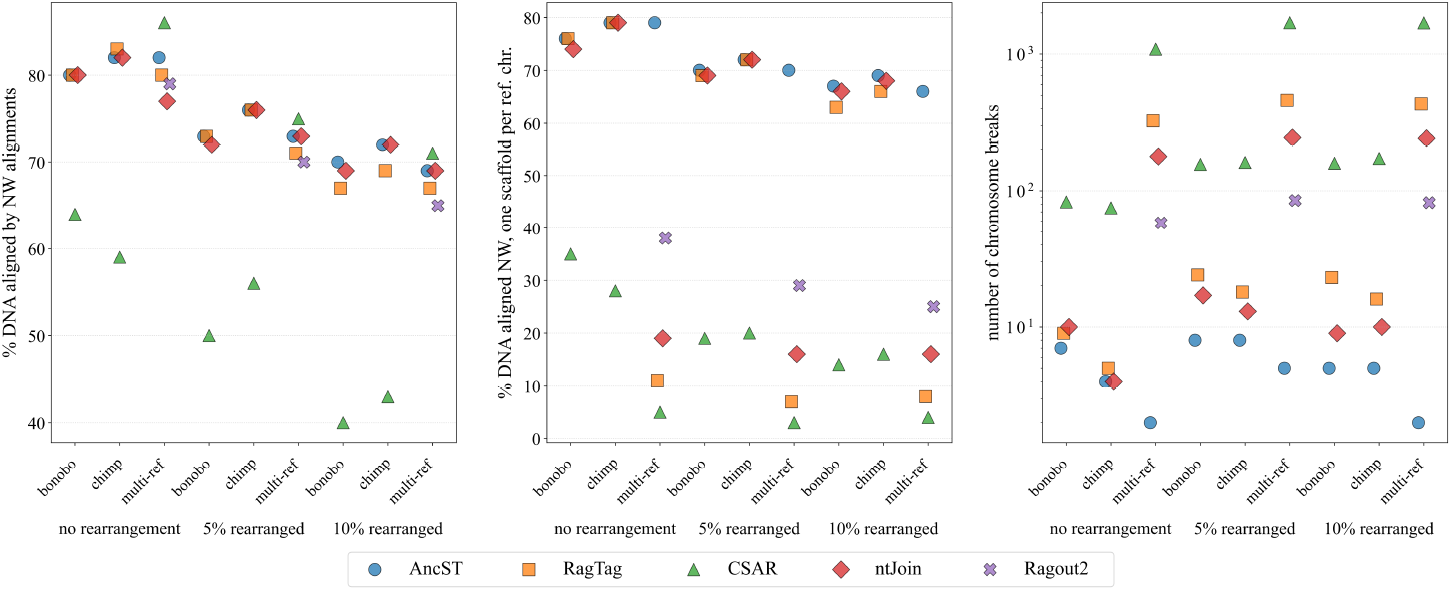
Key measures evaluating scaffolding performance for human synthetic fragmentation experiments on all tools. On the x-axis the genome version used as a target is listed below tick labels and three respective columns are plotted for each of them which represent the references used. Horizontal jitter is applied to data points within x-axis tick boundaries to separate overlapping markers. **Left:** % DNA aligned by the NW alignments of the new scaffolds; **Middle**: % DNA aligned by the NW alignments only allowing one new scaffold per reference chromosome; **Right:** the number of chromosome breaks on a log scale.

In the resulting perfect matching, head and tail of the same contigs are connected, resulting in a regular graph Y of vertex degree 2, i.e., a vertex disjoint union of cycles. After breaking each cycle by removing its minimum weight edge and removing completely unsupported edges with weight 0, we are left with a disjoint union of paths, each of which specifies a linear order of contigs as well as the reading direction of each contig, and thus a scaffold as depicted in Figure 1.

### 2.4 Synthetic Test Data

We produced three simulated draft assemblies of the human genome (GRCh38/hg38) in which the genome is chopped at repetitive sequences to generate contigs, based on typical fail scenarios of common assemblers where sequences cannot be uniquely elongated. In addition to this merely fragmented version we produced two further scenarios with additional noise (rearrangements) using a combination of tandem duplications, translocations and inversions; see Suppl. Sect. A.1 which can be found at Zenodo. The genome assemblies of chimpanzee (Clint PTRv2/panTro6) and bonobo (Mhudiblu PPA v0/panPan) were used as references for scaffolding. All sequence data were downloaded from the UCSC genome browser^d^.

### 2.5 Real-life Test Data

In addition to synthetic data, we complement benchmarks with real-life datasets. We employ our scaffolding pipeline to a set of 20 lower-quality `Drosophila` genomes (labeled as `scaffold` or `contig` assembly) published on NCBI, for 11 of which chromosome-level high-quality genomes are also available as references (Suppl. Tbl. 3). Here we focus on the application scenario in which genomes of multiple related species of limited assembly quality are available for homology-based scaffolding.

### 2.6 Scaffolding

We compared our approach with some widely used homology-based scaffolding approaches: `CSAR` ^3^ and `Multi-CSAR` ^8^, `RagTag` ^7^ with their `merge` command for multi-ref scaffolding, `ntJoin` ^5^ which allows multi-ref scaffolding as a default and `Ragout2` ^6^ which uses `cactus` multiple sequence alignments (MSAs) as input. The details about versions and parameters can be found in Suppl. Sect. C.

Supplementary Table 4 summarizes the algorithms used by each tool employed with additional references. To summarize, while `ntJoin` uses minimizers and `Ragout2` an MSA, `RagTag` and `CSAR` use sequence alignments to first obtain genetic marker regions which are shared across genomes. `AncST` uses a fully unique homology detection strategy, although some components outside of this anchor generation are derived from other tools. Duplications and otherwise repetitive sequences are filtered in a dedicated step in all tools except for `AncST` where those are excluded by design. For single-ref scaffolding `AncST` and `RagTag` share a conceptually similar greedy approach based on alignments while `CSAR` uses algebraic rearrangements. These three tools employ single-ref scaffolding before submitting the pairwise results to the same or similar graph-based optimization procedures in order to obtain multi-ref scaffolds. `ntJoin` employs its graph-based algorithms both for single- or multi-ref scaffolding, while `Ragout2` only performs multi-ref scaffolding. Thus, our implementation essentially uses a new type of genetic marker – synteny anchors – which is used in a greedy fashion relatively similar to the approach in `RagTag` for single-ref scaffolding. These pairwisely derived scaffolds are then reconciled using the same approach as in `Multi-CSAR`. The workflow of `AncST` is summarized schematically in Figure 1.

`AncST` was used to produce pairwise alignments between the respective genomes (parameters noted in Suppl. Sect. C.2). The resulting pairwise scaffolds were reconciled using a reimplementation of the ideas behind the merging algorithm in `Multi-CSAR` ^8^ as described in the respective *Merging of Alternative Scaffolds* section and further details can also be found in Suppl. Sect. C.2. While `Multi-CSAR` calculates weights for each reference; `RagTag`, `ntJoin` and `Ragout2` do not produce weights. Since weights may have a significant impact on the results, however, we ran the multi-ref experiment with `ntJoin` and `RagTag` of scaffolding the chopped but not further rearranged version of the synthetic human genome with the weights used by our pipeline and noted that the results were exactly the same as for unit weights. Furthermore, for experiments with real data we show in Suppl. Sect. B.4 that using weights from our pipeline in `RagTag` and `ntJoin` does not significantly alter the results. `Ragout2` does not use weights but an MSA produced by `cactus` for which a weighted initial phylogeny can be provided.

### 2.7 Evaluation

As previously described, contiguity measures alone are too simplistic to properly capture assembly quality ^14,15,16,17^. Hence, we employ several custom measures for a more comprehensive study of the resulting scaffolds’ qualities. Firstly, we use a Needleman-Wunsch global alignment algorithm ^18^ to quantify the edit distance between reconstructed scaffolds and the correct order of the contigs in the reference. A score of 1 is used for matching contigs, while insertions, deletions, and mismatches are scored with 0. The total score of the optimal alignment therefore equals the number of matched contigs for an optimal solution and, after determining the correct orientations of the aligned contigs, can directly be used as a performance measure. More precisely, we compute the *precision* as the percentage of the Needleman-Wunsch alignment score divided by the maximally achievable score, which is given by the total number of contigs minus 1/2 times the number of fused contigs. Alternatively, we may normalize the Needleman-Wunsch alignment score by the total number of contigs that are placed into any scaffold, resulting in a measure that is akin to *specificity*. Throughout, we refer to these measures as *NW prec*. and *NW spec*., respectively.

We also compute the total length of DNA covered by the correctly aligned and oriented contigs, i.e. exactly those with positive contribution in the NW alignment, normalized by the total length of all contigs from the original genomes (abbreviated as *NW DNA cov*.).

In addition to evaluating the alignments, we count the number of correctly connected contigs among all possible correct connections (*precision*) and all made connections (*specificity*). Similar to NW, we also compute the total length of DNA covered by correctly connected contigs relative to genomic length. Here, exactly all contigs are included that are connected correctly on at least one side, even if they are also involved in a false connection. We further refer to this metric as *adj. DNA cov*. Finally, we record the number of *breaks* which are introduced to the original chromosomes when they are covered by more than one newly assembled scaffold.

Finally, we employ the widely used assembly evaluation tool `Quast` (`v5.3.0`) ^19^ and compare the contiguity statistic *auN* (`a`rea `u`nder `N`x curve, see ^20^) for which a higher value indicates better performance. Additionally, we consider the number of *misassemblies* (for which a low number is preferable) reported against the respective reference genomes for both the synthetic and real datasets. We employ `auNA` as a measure of contiguity which takes into consideration only correct alignment blocks with the reference genome.

In summary, we highlight the global correctness of assemblies by NW alignments, chromosome breaks and `auN/A`. The number of misassemblies is computed by `Quast` if references are available. Local contig connections are assessed by the adjacency statistics.

## 3. Results

### 3.1 Synthetic Data – Custom Metrics

Table 1 summarizes the performance results for the synthetic human test data without adding additional noise. Supplementary Tables 1 and 2 in Suppl. Sect. A.4 show the results for the rearranged assemblies. Across the benchmark suite, no single tool dominates all measures, with different approaches showing advantages for different measures depending on the scenario. Figure 2 summarizes the key performance measures of our benchmarks. Percent of DNA covered by the NW alignments and the number of chromosome breaks arguably demonstrate the overall usefulness of the resulting scaffolds best. `AncST` consistently produces seemingly realistic single- and multi-ref scaffolds across all rearrangement levels. At the same time, `AncST` produces the fewest chromosome breaks across all experiments and scores the best or within four percentage points of the best performing tool for both NW DNA aligned measures. Considering the high number of chromosome breaks in all other tools regarding multi-ref scaffolds, ranging between 58 and 1692, it can be argued that `AncST` performs overall best with at most 5 breaks. Moreover, the number of chromosome breaks reduces for `AncST` in multi-ref scaffolding while it increased for all other tools compared to single-ref scaffolding. Even `Ragout2`, which was developed for multi-ref scaffolding, produces numerous chromosome breaks and does not cover more than 38 % DNA when only aligning the best matching scaffold per reference chromosome. In contrast, `AncST` produces scaffolds covering at minimum 66 %. Single-ref `CSAR` and multi-ref `Ragout2` generally perform best for the adjacency statistics, with values over 90 for the fragmented human genome but suffer, much as the other two tools, from many chromosome breaks. They thus produce correct local connections but fail to assemble them to larger pseudomolecules. In addition to single- and multi-ref `AncST`, only single-ref `ntJoin` and `RagTag` runs produce large pseudomolecules at chromosome break numbers of four to 24 and NW statistics which differ at 10 percentage points at most between those tools and respective experiments. Keeping in mind that our reconciliation algorithm is the same as in `Multi-CSAR` and similar to `RagTag`, with which we in turn share a similar greedy single-ref algorithm for alignments, this advantage highlights that improvements can be traced back in large parts to our unique anchor strategy. These results suggest that `AncST` anchors provide a more robust signal for multi-ref scaffold reconciliation compared to classic pairwise alignments, minimizers, algebraic rearrangements with pairwise alignments, or even multiple sequence alignments.

**Table 1.** Comparison of scaffolding performance. Data refer to the synthetic contig sets obtained by fragmenting the human genome. The genomes of chimpanzee (tro) and bonobo (pan) are used as references. We refer to Section 2.7 for details on these measures. The second columns for the alignment-based statistics in brackets considers the case in which only one new scaffold is allowed to be aligned to each reference chromosome as opposed to the figure before the brackets for which multiple scaffolds can align to each reference chromosome. All values (except for time measurements and the number of chromosome breaks) are percentages. Some of the best and worst performances are highlighted for each column (values in brackets are considered as separate columns except for the second time one). Accordingly, red means worst performing, magenta is within 3 percentage/time/count points of the worst, blue means best performing and cyan is within 3 percentage/time/count points of the best. Execution times are approximate wallclock times on a Linux server with 256 GB RAM and 64 available threads on two Intel(R) Xeon(R) Gold 6130 CPU @ 2.10GHz processors. The second time column reports the running time of (1) `AncST` without the pre-computation of anchors for each genome, since the latter can be performed as a one-time preprocessing for each genome and (2) `cactus` for `Ragout2`.

| Pipeline | Time[min] | NW |  |  | adj. |  |  | chr. breaks |
| --- | --- | --- | --- | --- | --- | --- | --- | --- |
|  |  | prec. | spec. | DNA cov. | prec. | spec. | DNA cov. |  |
| CSAR pan | 3200 | 65 (33) | **63** | 64 (35) | **94** | 90 | **91** | 83 |
| CSAR tro | 3200 | **63** (31) | **62** | **59** (28) | **95** | 91 | **91** | 75 |
| Multi-CSAR | **6500** | **87** (3) | 82 | **86** (5) | **43** | **68** | **70** | **1086** |
| RagTag pan | 25 | 79 (**77**) | 82 | 80 (**76**) | **93** | 90 | 87 | 9 |
| RagTag tro | 25 | 82 (**80**) | 84 | **83** (**79**) | **92** | 89 | 86 | **5** |
| Multi-RagTag | 52 | **60** (9) | 92 | 80 (11) | 48 | 86 | 75 | 327 |
| ntJoin pan | **20** | 79 (**77**) | 81 | 80 (74) | **92** | 88 | 82 | 10 |
| ntJoin tro | **20** | 82 (**80**) | 83 | 82 (**79**) | **92** | 87 | 82 | **4** |
| Multi-ntJoin | 28 | **89** (28) | **96** | 77 (19) | 85 | 93 | 74 | 178 |
| Ragout2 (multi) | 4100 (15) | 74 (39) | 80 | 79 (38) | **93** | **97** | **91** | 58 |
| AncST pan | 600 (14) | 79 (76) | 84 | 80 (**76**) | 91 | 91 | **88** | 7 |
| AncST tro | 600 (14) | 82 (**79**) | 87 | 82 (**79**) | **92** | 92 | **88** | **4** |
| Multi-AncST | 1200 (30) | 81 (**79**) | 86 | 82 (**79**) | **92** | 91 | **88** | **2** |

To assess the robustness of the different tools under assembly noise, here simulated as rearrangements in synthetic human data, we plot the absolute relative changes of all performance measures from Table 1 when changing from the fragmented version of the human genome to either of the two rearranged ones as shown in Figure 3 on the left. `ntJoin` and `AncST` show the lowest variability between the fragmented and rearranged versions of the human genome. Similarly, on the right we plot the absolute relative changes of performance measures when using the different reference genomes including using both. In accordance with the results presented above, `AncST` is much more robust to using multiple references, thus showing the lowest variability by far. In Suppl. Sect. C.4 we show that also when only considering differences between using bonobo or chimp as references `AncST` is arguably the most robust.

**Fig. 3.**
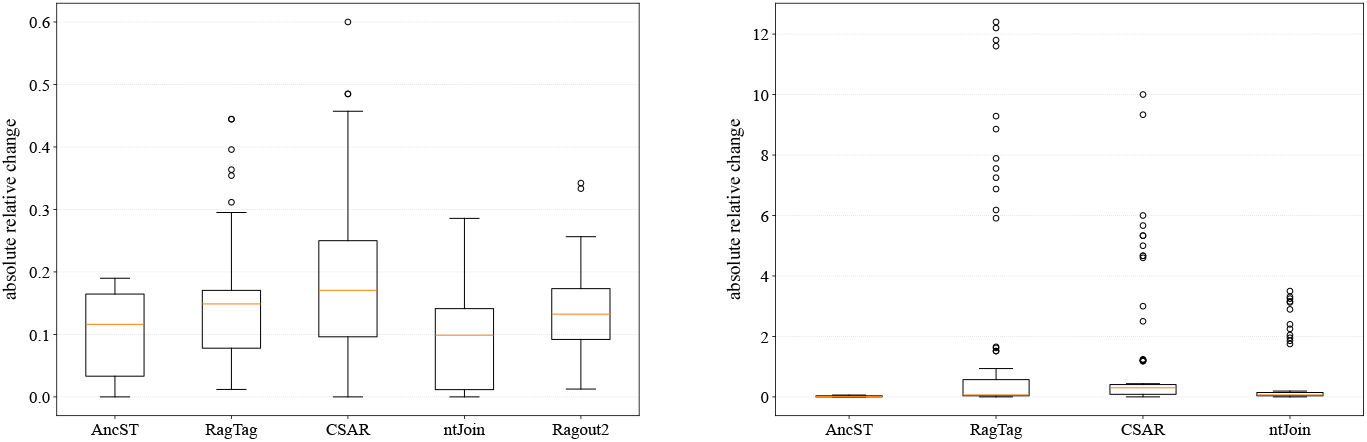
Boxes span the interquartile range (Q1-Q3); the horizontal line marks the median. Whiskers extend to the most extreme data point within 1.5 *×* IQR of the box; values beyond are shown as individual points (outliers). On the left, the absolute relative changes for all performance measures from Table 1 are shown between the experiments with the fragmented version of the human genome and the two additionally rearranged ones (18 data points for `Ragout2`: 9 performance measures *×* 2 rearrangement scenarios; 54 data points for the other tools: 3 different references *×* 9 performance measures *×* 2 rearrangement scenarios). On the right the relative changes are shown between using either or both of the references (81 data points: 3 ref comparisons *×* 9 performance measures *×* 3 human genome versions).

In terms of speed, the minimizer-based `RagTag` and `ntJoin` are orders of magnitude faster than the other tools if the extensive alignments needed by the other tools are not precomputed. A more detailed breakdown of the resource consumption of the `AncST` pipeline shows that about a quarter of the compute time is used for the collection of the k-mer statistics with `GenMap` ^21^ as a basis for determining the anchor candidates. Further two thirds of the effort is expended for the computation of *d*_0_-uniqueness of these candidates through `blast` comparisons of the anchor candidates against their own genomes. All other tasks in the pipeline account for less than 10 % of the wallclock time. Hence, the computation time for pairwise anchor candidate comparison and scaffolding alone are on par with those of the fastest methods.

### 3.2 Synthetic Data – Evaluation with Quast

`Quast` evaluation further supports the results obtained with the measures described above. The full results and corresponding figures are compiled in Suppl. Sect. A. In summary, `CSAR` performs worst overall, with one exception: It shows the best `auNA` metric for single-ref scaffolding of the chopped but not rearranged version of the human genome. In comparison with the NW scores in Table 1 this outlier suggests that there may only be few misassemblies preventing a higher NW score since original contigs are not allowed to align twice while `auNA` considers sequence alignments and can thus break new scaffolds. For the multi-ref scenario AncST shows robustness while the other tools can usually not only *not* benefit from using multiple references but performance drops significantly except for `ntJoin` with regard to `auNA`.

In Suppl. Sect. A.3 we plot the relative coverage of the 24 human chromosomes (22+`X`+`Y`) estimated by the total length of all `Quast` alignments with the new scaffolds covering this chromosome. It can be seen that while for single-ref homology-based scaffolding all tools except `CSAR` produce sensibly large scaffolds, only our pipeline achieves this for multi-ref runs.

### 3.3 Real-life Data – Evaluation with Quast

As for the synthetic data above, we compare `auN`, `auNA` and the number of `misassemblies` as computed by `Quast` in Figure 4 and suppl. Sect. B.2. We excluded `Multi-CSAR` from the experiments as it implies resource-intensive whole-genome alignments of all pairs and this tool shows largely disappointing results for the synthetic dataset. In In Figure 4 it can be seen that the tools are relatively comparable in their results but `AncST` performs best most consistently.

**Fig. 4.**
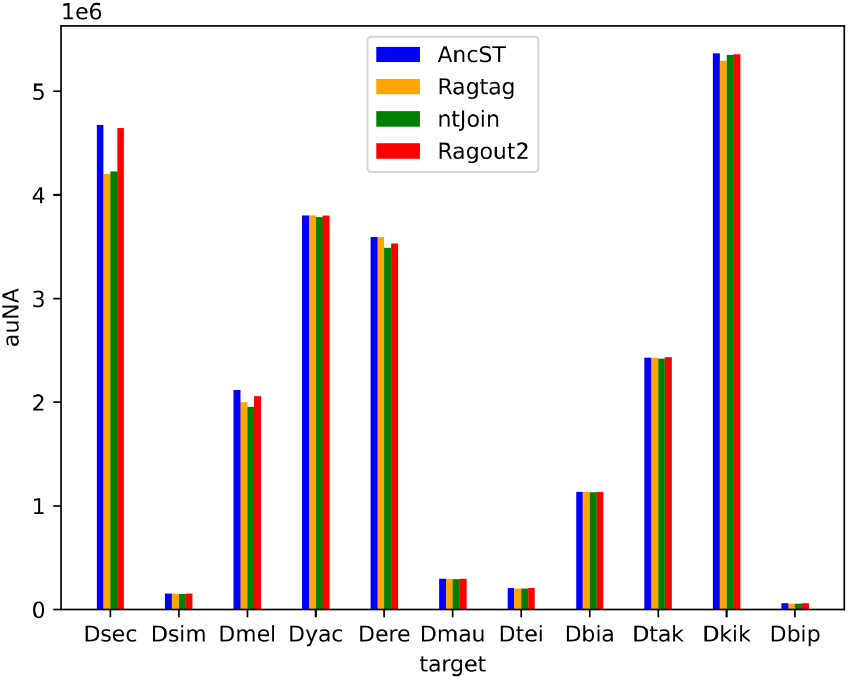
`auNA` as computed by `Quast` for the 11 `Drosophila` newly scaffolded species with a chromosome-scale reference genome on NCBI. Further details about the species in Suppl. Tbl. 3.

In the following, the detailed analyses are restricted to the two most promising tools based on the contiguity statistics `auN` and `auNA` which are our own and `RagTag`. Firstly, in order to assess whether the multi-ref approach improves over single-ref scenarios, as one would expect from the low quality of the genomes, we compare the fractions of single-ref scaffolds with higher contiguity as measured by `auN`. Comparing all 380 single-ref scaffolds with their respective multi-ref version, in 76 % of cases from `RagTag` the multi-ref version is more contiguous while our pipeline achieves higher contiguity in 90 % of cases. Hence, while both pipelines can utilize the multi-ref approach for a scenario in which a set of lower-quality genomes is available for scaffolding, our pipeline more consistently produces multi-ref scaffolds of higher contiguity. To estimate how many unrealistically large scaffolds are created in either approach, we count the number of cases in which the largest scaffold exceeds 50 million nts, a number significantly higher than any `Drosophila` reference chromosome. While no single-ref scaffold exceeds 50 million nts for either approach, `RagTag` produces 8 and `AncST` 4 multi-ref scaffolds longer than this threshold. Thus, both approaches produce unrealistically large scaffolds in a significant number of cases of the multi-ref approach but the issue is more severe for `RagTag`. To further investigate the scaffolds’ contiguity and severity of potential misassemblies we compare the coverage of large chromosomes of the reference genomes (if available) with potential new scaffolds and single original contigs in some cases. The complete results for each species are displayed in Suppl. Sect. B.3. As an example, consider the extreme case of `Drosophila biarmipes` whose `RagTag` scaffolds display an `auN` value of around 176 million and a longest contig of length 180 million while the total assembly length is 185 million. Related to the proportion of unrealistically large scaffolds, `RagTag` scaffolds show more cases in which multiple reference chromosomes are covered by the same scaffold. In order to assess whether `RagTag`’s performance can be improved by using weights for each reference we repeated the scaffolding and analysis using the same weights as in our pipeline. In essence, using our weights for `RagTag` does not significantly improve its performance (Suppl. Sect. B.4) for this multi-ref scaffolding experiment.

## 4. Discussion

Given the NP-completeness of the problem as well as large datasets and code bases employed, we can only speculate about the algorithmic reasons for specific differences in performance measures.

Theoretically, whole-genome alignments from `MUMmer` ^22^ in `CSAR` and `cactus` for `Ragout2` should achieve the highest resolution. The performance benchmarks show, however, that `CSAR` and `Multi-CSAR` arguably perform the worst overall and `Ragout2` only excels in contig adjacencies while producing relatively fragmented scaffolds. We suspect that these tools do not resolve the conflicting information from duplicated and rearranged sequences with sufficient accuracy and that these problems then propagate to the scaffolding algorithms. The algebraic rearrangement models implemented in `CSAR` do not seem to capture the important signals for single-ref scaffolding very well with consistently worst performance measures with few exceptions as described above. Hence, this algorithm can capture correct local contig connections but misses too many pieces to produce sensible pseudomolecules and additionally produces apparently conflicting single-ref scaffolds. Moreover, `Multi-CSAR`’s implementation of the maximum perfect matching algorithm may be suboptimal because we observe similar or better results for the multi-ref scaffolds than for its single-ref inputs with our implementation.

Except for `CSAR`, all other tools perform acceptably and comparable in the single-ref scenarios exemplified by the DNA coverage of NW alignments which range narrowly between 80 and 83 %. In the multi-ref scenarios, only `Ragout2` and `AncST` perform acceptably when looking at key measures for the synthetic data experiments: All other tools produce at least twice as many chromosome breaks and cover at most half the DNA in the NW alignments where only one scaffold is allowed to align. Thus, the other tools produce substantially smaller pseudomolecules.

`Ragout2` uses several sophisticated steps to first obtain synteny blocks of different granularity from the MSA and then refine the breakpoint graph. In contrast, our implementation only makes use of atomic synteny anchors without producing larger blocks first. Despite its sophisticated algorithms, `Ragout2` performs worse than `AncST` except for correct contig adjacencies. Most likely, our anchors already provide a robust signal for scaffolding without further refinement of synteny blocks. In fact, we confirmed that using `MCScanX` ^23^ collinearity blocks instead of single anchors did not improve the results of `AncST`. We speculate that the superior performance of `AncST` in both synthetic and real-world multi-ref settings could be due to the finer granularity and overlap characteristics of anchor matches. While `AncST` typically produces anchors of hundreds to thousands of nts, `Ragout2` uses initial synteny blocks of few hundred nts and `ntJoin` uses a minimizer for 1000 nts long windows, `CSAR` and `RagTag` use “longest” alignments which could be at least several thousand nts long. Thus, this interpretation is reconcilable with the latter two performing even worse in most performance measures than the former. Moreover, `AncST` anchor candidate boundaries are the same for each pairwise comparison resulting in implicit multiple sequence matches which may introduce fewer conflicting contig connections for multi-ref scaffolding. Accordingly, we observe that the number of cycles broken to obtain scaffolds from the maximum perfect matching is extremely small in our pipeline, which eventually leads to the more contiguous scaffolds. This should also apply to `ntJoin` and `Ragout2` which use minimizers present in all references and an MSA, respectively. Both perform better in most measures contrasting `CSAR` and `RagTag` which rely on pairwise alignments with their pair-specific boundaries. Hence, perhaps `AncST` outperforms `ntJoin` and `Ragout2` for multi-ref scaffolding because it does not require matches to be present in all references but allows information from pairwise alignments to be incorporated.

Since only closely related species are used for homology-based scaffolding, `AncST`’s phylogenetic performance boundaries ^10^ are of no practical concern. Other factors potentially limiting performance, such as reference assembly quality, repeat content or polyploidy, theoretically apply to all tools. Whether polyploid genomes can effectively be aligned by `AncST` depends on the relative timing of chromosome duplications: if they are sufficiently old, enough duplicated sequences may have diverged to serve as anchor candidates. A major drawback of our pipeline is the run time and possibly high demands on memory for computation of the string indices which may be prohibitive in the case of very large genomes. In this respect, it cannot compete with `RagTag` and `ntJoin`, except if anchors are precomputed, while being more scalable than `CSAR` and `Ragout2` avoiding a full whole-genome alignment. Since we develop `AncST` as a synteny-based comparative genomics toolbox and provide a webserver for anchor computation and download of precomputed ones, our scaffolder can be accessed relatively easily despite the possible resource and other practical challenges.

## 5. Conclusions

Fragmented assemblies systematically inflate apparent gene counts and produce truncated or split gene models that propagate into downstream annotation ^24,25^, and they limit the resolution of any analysis that depends on long-range coordinates, such as TAD analyses. Scaffolding can alleviate such issues ^14,15,16^, and homology-based scaffolding is likely to gain increasing importance in non-model-organism genomics in response to the rapidly growing number of high-quality genomes that are suitable as references. The phylogenetic distance of available reference genomes to a newly sequenced genome thus shrinks, increasing the accuracy of the reference-based approach and making multi-ref scaffolding more attractive. A single-ref scaffold inevitably encodes the rearrangement state of that one reference, so rearrangements that are species-specific to the chosen reference are imprinted onto the target as pseudo-rearrangements ^26,27^. Hence, the methodology has the potential to provide biologists with large numbers of accessible (close to) chromosome-level genome assemblies for functional analyses, comparative genomics, and evolutionary studies at moderate experimental and computational costs. This is of particular relevance with respect to the large-scale efforts mentioned in the introduction.

We have shown that anchor candidates can be used effectively for homology-based contig scaffolding. Our anchor-based approach occupies a middle ground in the scaffolding landscape: computational cost and scalability fall between expensive whole-genome alignment methods and highly efficient minimizer-based approaches, while achieving comparable accuracy to established tools.

For single-ref scaffolding `RagTag` and `ntJoin` are clearly the best performing tools overall, particularly considering their extremely low run time. Hence, minimizer sketches and alignments based on them seem to provide most of the relevant signal for homology-based scaffolding and even avoid some of the issues leading to inferior performance of `CSAR` and `AncST`. Nevertheless, `AncST` still achieves comparable performance in single-ref scenarios since its NW alignments cover nearly as much DNA as the best performing tools, the adjacency statistics are within a few points of competing tools, and `AncST` produces the smallest number of chromosome breaks. For multi-ref scenarios, `AncST` outperforms all other tools with `Ragout2` performing second best. By reconciling evidence across multiple references, `AncST` produces a less reference-biased ordering, since rearrangement events specific to any individual reference receive less weight in the consensus. This is also reflected empirically in our robustness analysis (Fig. 3, right), where `AncST` shows by far the lowest variability across reference choices.

Improvements in structural assembly quality have direct consequences for biological inference. Many comparative genomics methodologies rely on accurate contiguous assemblies. For example, reference-based gene annotation and annotation lift-over pipelines such as `Liftoff` ^28^ and `TOGA` ^29^ explicitly assume that contigs are correctly ordered and oriented, and break or mis-call genes when they are not. Macrosynteny analyses, e.g. the reconstruction of ancestral linkage groups ^30,31^, require chromosome-scale scaffolds to resolve gene-order conservation across hundreds of millions of years. Likewise, structural-variant calling depends on chromosome-scale inputs as large rearrangements are unresolvable on fragmented assemblies ^32,33^.

The issues of homology-based scaffolding are inherently addressed in the nature of `AncST` anchors: By design, repetitive sequences are excluded and ambiguous alignments of anchors are marked as unreliable. This design may also facilitate the overall superior performance of `AncST` with respect to the real data experiments where we scaffolded explicitly poor assemblies with higher content of repeats and misassemblies. Furthermore, anchor candidates are computed for highly fragmented assemblies in just the same way and can readily indicate problematic genomic regions when their alignments to other genomes’ candidates are ambiguous. As these alignments are marked as unreliable and not further used, performance can stay relatively high even for poor assemblies. A distinctive advantage of the anchor-based approach is that candidates can be precomputed and stored as simple annotation files that represent anchors as coordinate pairs associated with their level of the interval’s *d*_0_-uniqueness. Consequently, anchors of the target genome are the same in each pairwise comparison with the references which may promote the observed robustness for reconciliation of multiple single-ref scaffolds.

A natural next step is leveraging non-collinear anchor alignments to identify breakpoints arising from genome rearrangements. The anchor matches themselves, moreover, can be fed directly into the analysis of macrosynteny, which is usually performed as part of genome assembly projects. For instance, we can readily track possible inversion scenarios by enumerating regions for which `AncST` anchors align on the opposing strand in considerable numbers. Translocations are indicated by significant collinear stretches of alignments with a breakpoint in one of the species. The current implementation of the `AncST`-based pipeline identifies contigs with alignments to more than one reference chromosome and those with alignments in opposite reading directions on the same reference chromosome. These are strong indicators for rearrangements relative to the reference genome. In principle, these data could be used to extend scaffolds beyond such breakpoints. Whether this can be done with acceptable reliability, however, is currently subject to investigation.

## Acknowledgements

This work was funded by the German Research Foundation as part of SPP 2349 *“Genomic Basis of Evolutionary Innovations (GEvol)”* (grant no. 502862570). PFS acknowledges the financial support by the Federal Ministry of Education and Research of Germany (BMBF) through DAAD project 57616814 (SECAI, School of Embedded Composite AI), and jointly with the Sächsische Staatsministerium für Wissenschaft, Kultur und Tourismus in the programme Center of Excellence for AI-research *Center for Scalable Data Analytics and Artificial Intelligence Dresden/Leipzig*, project identification number: SCADS24B.

## Availability and Supplementary Material

The appendix and AncST anchors for the human dataset are available at Zenodo (https://zenodo.org/records/21793010). Further supplementary material is available at https://zenodo.org/records/15061775. The prototypical pipeline is included in the supplementary material and `AncST` Github repository (https://github.com/Norsbus/AncST) as well as part of results from the webserver (https://anchored.bioinf.uni-leipzig.de/).

## Footnotes

a https://www.darwintreeoflife.org

b https://www.erga-biodiversity.eu

c https://www.earthbiogenome.org

d https://hgdownload.soe.ucsc.edu/goldenPath/

