## Appendix for "Anchors for Homology-Based Scaffolding"

### A Benchmark of Synthetic Human Data

#### A.1 Simulation of Synthetic Fragmentation

Locations of rearrangements were restricted to respecting annotation items, i.e. breakpoints were only introduced at the start or end of annotated items in the annotation file of the human genome. We distinguish between short-range and long-range rearrangements. The former only affect a single annotation item, while the latter pertain to an interval containing two or more annotation items separated by not more than 1 million bases. In addition to a local inversion of the interval, rearrangements were defined as follows: For translocation we randomly selected a different chromosome, an insertion position, and the orientation of the inserted fragment. Up to four duplication events are allowed, either local or remote. For a local duplication the copy of a selected interval is inserted, possibly in inverted orientation, at a location between 1000 and 100000 bases downstream of the interval in question. Alternatively, duplicates were inserted in a randomly chosen chromosome. If a translocated sequence was chosen to only be translocated but not copied, then the original source interval was deleted in order to avoid bloating the genome size. We created one version in which at least 5 % of the human genome are rearranged and another one in which at least 10 % are rearranged. The number of duplication events as well as sequence intervals are chosen arbitrarily. The rationale behind the moderate number of rearrangement events in the test data is that contigs with a large number of rearrangements relative to the reference cannot be safely placed into a scaffold since homology-based scaffolding only can be employed when co-linearity is the norm. The rearranged genomes are documented in the supplementary material. Contigs were extracted in the same manner from both the rearranged and conserved human genome. In order to mimic the fact that contig assembly usually stops at repetitive sequences, we defined contig ends at stretches of at least 10000 N in the hard-masked version of the human genome. In addition, we introduced random breakpoints within the initial fragments. At this stage we only accepted contigs consisting of at least 20000 bases and 30 % unmasked sequence. Each initial fragment was subjected to 30 repetitions of the random fragmentation attempts. In order to simulate assembly errors [1], we fused 5 % of the resulting fragments at random. Moreover, reading direction was determined at random.

#### A.2 auN(A) and Misassemblies as Computed by Quast

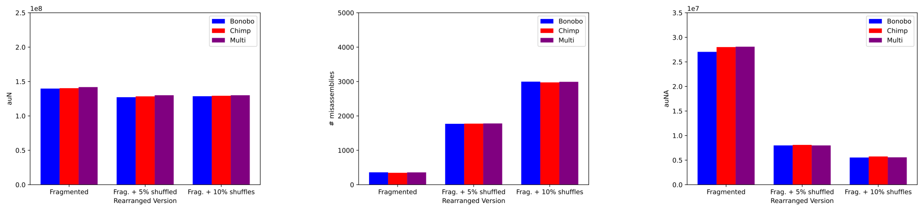


**Fig. 1** (Left:) *auN*, (Middle:) *misassemblies* and (Right:) *auNA* as computed by `Quast` for the new scaffolds produced by `AncST`.

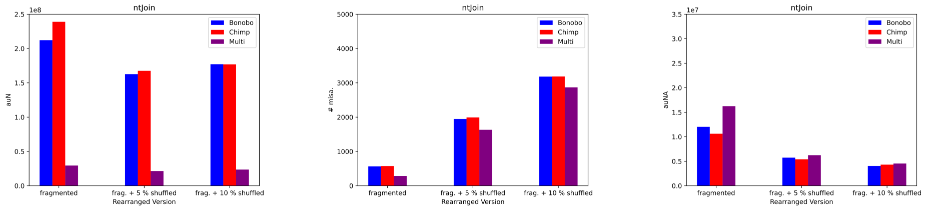


**Fig. 2** (Left:) *auN*, (Middle:) *misassemblies* and (Right:) *auNA* as computed by `Quast` for the new scaffolds produced by `ntJoin`.

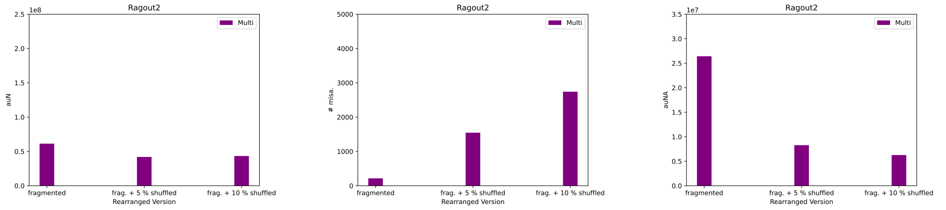


**Fig. 3** (Left:) *auN*, (Middle:) *misassemblies* and (Right:) *auNA* as computed by `Quast` for the new scaffolds produced by `Ragout2`.

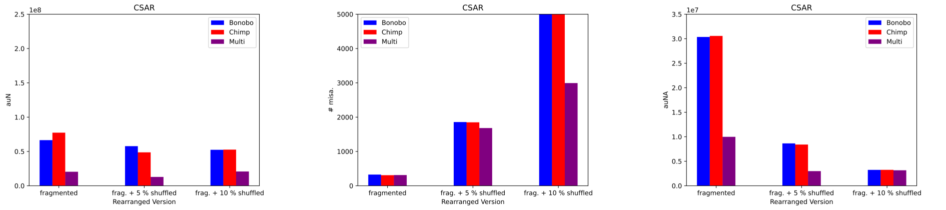


**Fig. 4** (Left:) *auN*, (Middle:) *misassemblies* and (Right:) *auNA* as computed by `Quast` for the new scaffolds produced by `CSAR`.

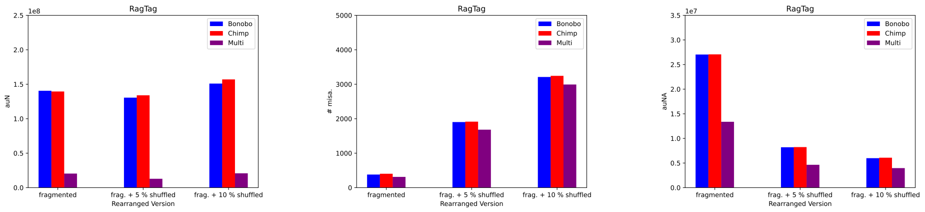


**Fig. 5** (Left:) *auN*, (Middle:) *misassemblies* and (Right:) *auNA* as computed by `Quast` for the new scaffolds produced by `RagTag`.

#### A.3 Coverage of Main Human Chromosomes with Quast Alignments of New Scaffolds

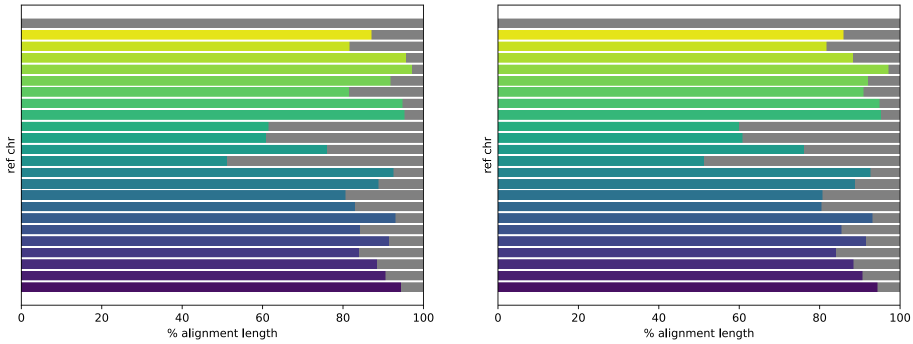


**Fig. 6** Shown are data based on the results of scaffolding the human genome with at least 5 % rearranged sequence against (left:) chimp and (right:) chimp and bonobo from `AncST`. Relative coverage (X-axis) of 24 human chromosomes (22+`X`+`Y`) (Y-axis) by all alignments produced by `Quast`. First, the total length of all alignments of a reference chromosome with any new scaffolds is noted. Then each new scaffold is assigned its relative coverage as the proportion of the length of its alignments with the reference. Scaffolds covering at least 50 % of a reference are colored and the rest is gray. Relative coverage by one contig is bordered by white vertical lines.

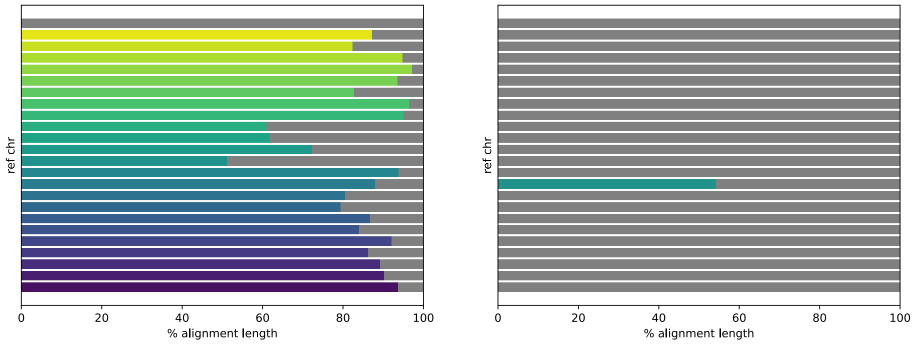


**Fig. 7** Shown are data based on the results of scaffolding the human genome with at least 5 % rearranged sequence against (left:) chimp and (right:) chimp and bonobo from `ntJoin`. Details can be found in caption of Fig. 6.

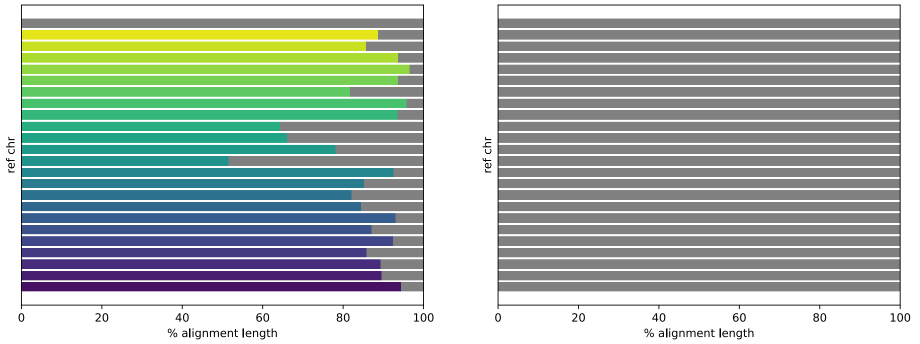


**Fig. 8** Shown are data based on the results of scaffolding the human genome with at least 5 % rearranged sequence against (left:) chimp and (right:) chimp and bonobo from `RagTag`. Details can be found in caption of Fig. 6.

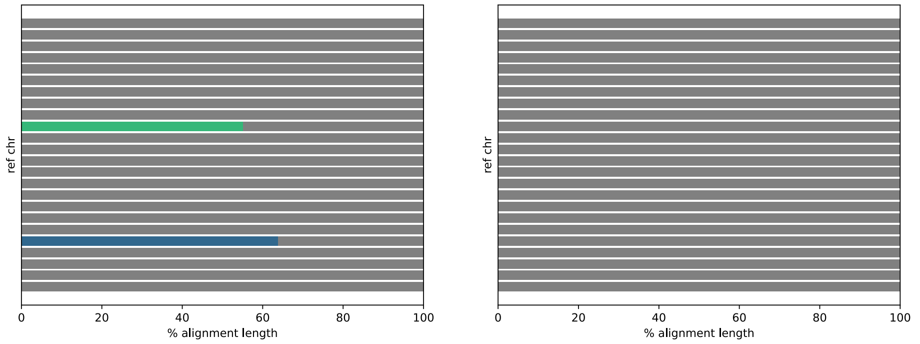


**Fig. 9** Shown are data based on the results of scaffolding the human genome with at least 5 % rearranged sequence against (left:) chimp and (right:) chimp and bonobo from `CSAR`. Details can be found in caption of Fig. 6.

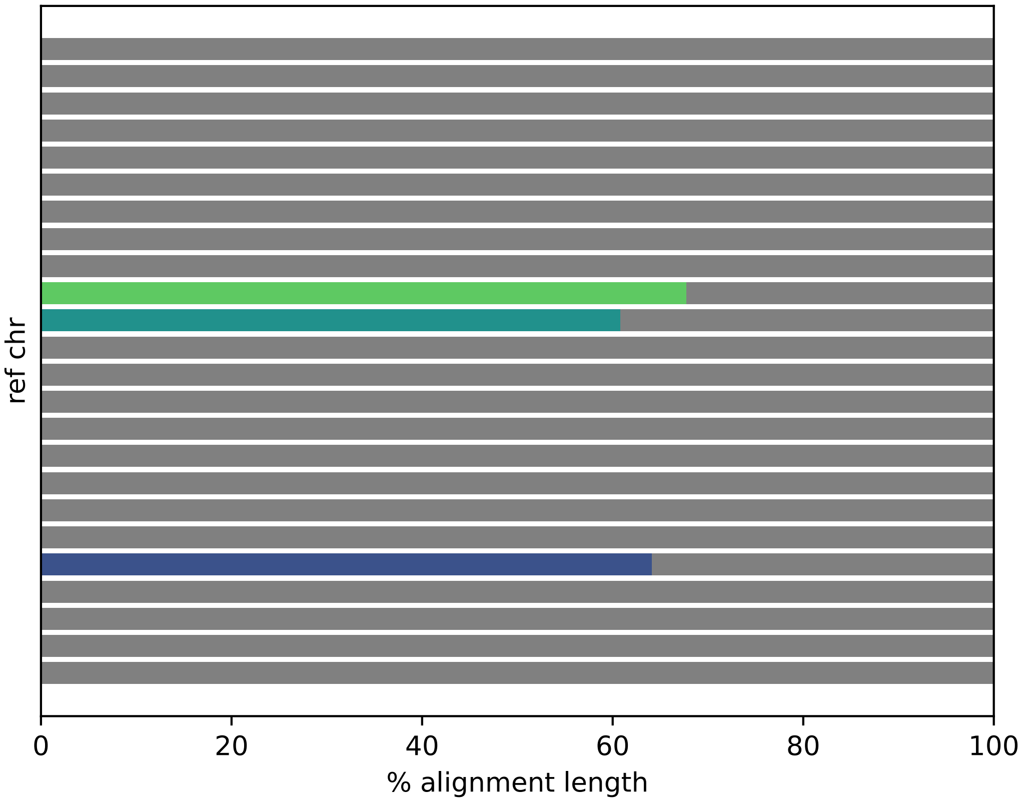


**Fig. 10** Shown are data based on the results of scaffolding the human genome with at least 5 % rearranged sequence against chimp and bonobo from `Ragout2`. Details can be found in caption of Fig. 6.

#### A.4 Summary of Benchmarks

**Table 1** Scaffolding results for the chopped human genome with at least 5 % rearranged sequence. The genomes of chimpanzee (tro) and bonobo (pan) are used as references. We refer to Section *Evaluation* for details on these metrics. The second columns for the alignment-based statistics in brackets considers the case in which only one new scaffold is allowed to be aligned to each reference chromosome as opposed to the figure before the brackets for which multiple scaffolds can align to each reference chromosome. All values (except for time measurements and the number of chromosome breaks) are percentages. Some of the best and worst performances are highlighted for each column (values in brackets are considered as separate columns except for the second time one). Accordingly, red means worst performing, magenta is within 3 percentage/time/count points of the worst, blue means best performing and cyan is within 3 percentage/time/count points of the best. Execution times are approximate wallclock times on a Linux server with 256 GB RAM and 64 available cores on two Intel(R) Xeon(R) Gold 6130 CPU @ 2.10GHz processors. The second time column reports the running time of (1) `AncST` without the pre-computation of anchors for each genome, since the latter can be performed as a one-time preprocessing for each genome and (2) `cactus` for `Ragout2`.

| Pipeline | Time[min] | NW | | | adj. | | | chr. breaks |
|---|---|---|---|---|---|---|---|---|
| | | prec. | spec. | DNA cov. | prec. | spec. | DNA cov. | |
| CSAR pan | 3200 | 50 (17) | 53 | 50 (19) | 82 | 85 | 83 | 156 |
| CSAR tro | 3200 | 53 (19) | 56 | 56 (20) | 83 | 83 | 85 | 161 |
| Multi-CSAR | 6500 | 79 (3) | 80 | 75 (3) | 37 | 63 | 60 | 1692 |
| RagTag pan | 28 | 67 (64) | 79 | 73 (69) | 78 | 86 | 79 | 24 |
| RagTag tro | 26 | 71 (67) | 83 | 76 (72) | 78 | 86 | 79 | 18 |
| Multi-RagTag | 56 | 43 (5) | 89 | 71 (7) | 31 | 78 | 65 | 460 |
| ntJoin pan | 20 | 68 (65) | 81 | 72 (69) | 79 | 89 | 79 | 17 |
| ntJoin tro | 20 | 72 (68) | 85 | 76 (72) | 79 | 88 | 78 | 13 |
| Multi-ntJoin | 28 | 77 (20) | 96 | 73 (16) | 74 | 93 | 69 | 246 |
| Ragout2 (multi) | 4700 (18) | 66 (26) | 83 | 70 (29) | 79 | 95 | 83 | 85 |
| AncST pan | 650 (14) | 65 (63) | 82 | 73 (70) | 76 | 90 | 79 | 8 |
| AncST tro | 650 (14) | 69 (66) | 87 | 76 (72) | 76 | 89 | 80 | 8 |
| Multi-AncST | 1300 (30) | 66 (64) | 82 | 73 (70) | 77 | 89 | 80 | 5 |

**Table 2** Scaffolding results for the chopped human genome with at least 10 % rearranged sequence. For details we refer to the caption of Table 1.

| Pipeline | Time[min] | NW | | | adj. | | | chr. breaks |
|---|---|---|---|---|---|---|---|---|
| | | prec. | spec. | DNA cov. | prec. | spec. | DNA cov. | |
| CSAR pan | 3300 | 48 (17) | 51 | 40 (14) | 72 | 75 | 68 | 159 |
| CSAR tro | 3400 | 52 (18) | 55 | 43 (16) | 73 | 76 | 69 | 172 |
| Multi-CSAR | 6800 | 77 (3) | 78 | 71 (4) | 33 | 56 | 49 | 1687 |
| RagTag pan | 29 | 66 (63) | 77 | 67 (63) | 76 | 83 | 71 | 23 |
| RagTag tro | 27 | 70 (66) | 80 | 69 (66) | 77 | 83 | 72 | 16 |
| Multi-RagTag | 58 | 42 (5) | 85 | 67 (8) | 29 | 73 | 57 | 433 |
| ntJoin pan | 20 | 67 (65) | 81 | 69 (66) | 78 | 88 | 74 | 9 |
| ntJoin tro | 20 | 70 (67) | 84 | 72 (68) | 79 | 88 | 74 | 10 |
| Multi-ntJoin | 28 | 77 (22) | 96 | 69 (16) | 73 | 91 | 62 | 243 |
| Ragout2 (multi) | 4800 (18) | 64 (29) | 81 | 65 (25) | 78 | 95 | 77 | 82 |
| AncST pan | 650 (15) | 66 (64) | 82 | 70 (67) | 73 | 89 | 73 | 5 |
| AncST tro | 650 (15) | 69 (66) | 86 | 72 (69) | 77 | 89 | 74 | 5 |
| Multi-AncST | 1300 (32) | 66 (64) | 82 | 69 (66) | 77 | 89 | 73 | 2 |

### B Benchmark on Drosophila Assemblies

#### B.1 Genomic References

**Table 3** Genomes of `Drosophila` species used. `same` in `NCBI Ref.`(erence) means that the Contig or Scaffold level assembly used as a scaffolding target is also marked as the reference chromosome for this species on NCBI.

| Accession | Species | Abbreviation | Assembly Level | NCBI Ref. | Identifier |
|---|---|---|---|---|---|
| GCA_018904445.1 | `D. sechellia` | Dsec | Scaffold | GCF_004382195.2 | A |
| GCA_039725655.1 | `D. simulans` | Dsim | Contig | GCF_016746395.2 | B |
| GCA_000778455.1 | `D. melanogaster` | Dmel | Contig | GCF_000001215.4 | C |
| GCA_018904385.1 | `D. yakuba` | Dyak | Contig | GCF_016746365.2 | D |
| GCA_018904525.1 | `D. erecta` | Dere | Scaffold | GCF_003286155.1 | E |
| GCA_018904475.1 | `D. mauritiana` | Dmau | Scaffold | GCF_004382145.1 | F |
| GCA_018903625.1 | `D. teissieri` | Dtei | Contig | GCF_016746235.2 | G |
| GCA_005876975.1 | `D. orena` | Dore | Contig | same | H |
| GCF_018153835.1 | `D. eugracilis` | Deug | Contig | same | I |
| GCA_018148935.1 | `D. biarmipes` | Dbia | Contig | GCF_025231255.1 | J |
| GCA_018152695.1 | `D. takahashii` | Dtak | Contig | GCF_030179915.1 | K |
| GCF_018152265.1 | `D. ficusphila` | Dfic | Contig | same | L |
| GCF_018152505.1 | `D. elegans` | Dele | Contig | same | M |
| GCF_018152115.1 | `D. rhopaloa` | Drho | Contig | same | N |
| GCA_008042655.1 | `D. burlai` | Dbur | Scaffold | same | O |
| GCA_018152535.1 | `D. kikkawai` | Dkik | Contig | GCF_030179895.1 | P |
| GCA_008042735.1 | `D. leontia` | Dleo | Scaffold | same | Q |
| GCA_021223765.1 | `D. bipectinata` | Dbip | Scaffold | GCF_030179905.1 | R |
| GCA_018153235.1 | `D. malerkotliana` | Dmal | Contig | same | S |
| GCA_018148915.1 | `D. ananassae` | Dana | Contig | same | T |

#### B.2 Scaffolding Evaluation of Drosophilas

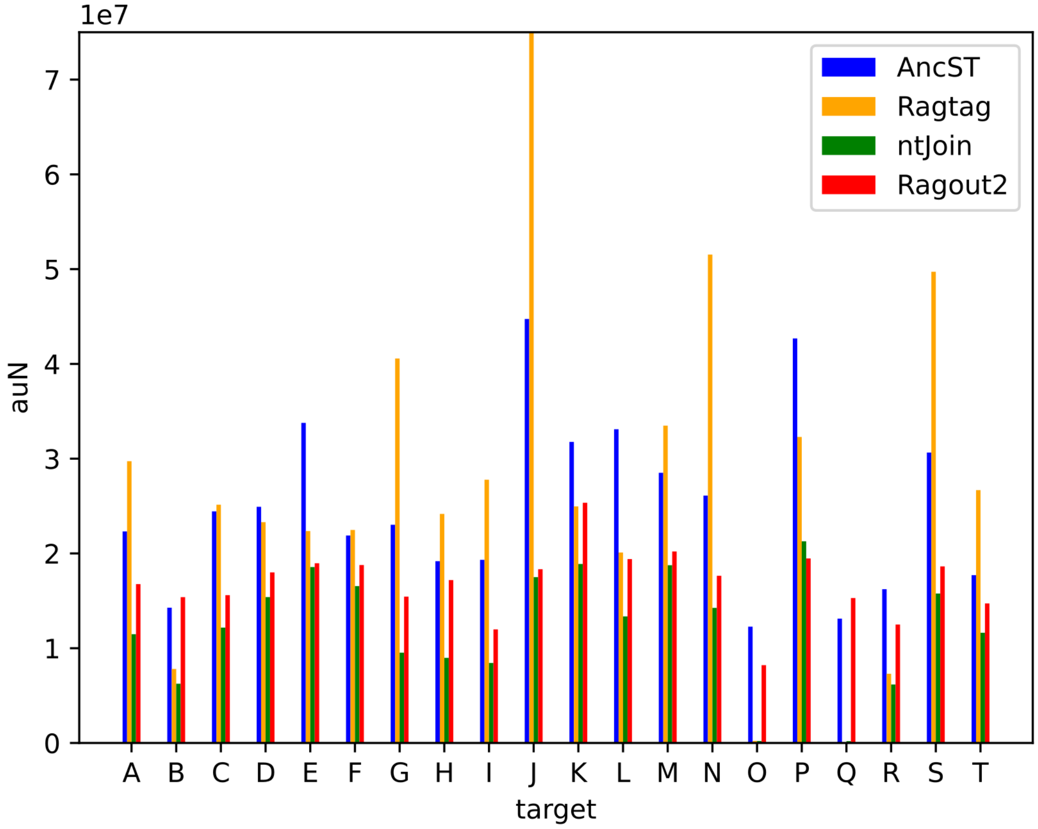


**Fig. 11** *auN* as computed by `Quast` for 20 `Drosophila` newly scaffolded species. The new scaffolds computed by `RagTag` for `Drosophila biarmipes` ($J$) show an *auN* of around 176 million while we set the upper limit of the y-axis to 75 million for clearer display. Identifier correspondence and further details in Table 3.

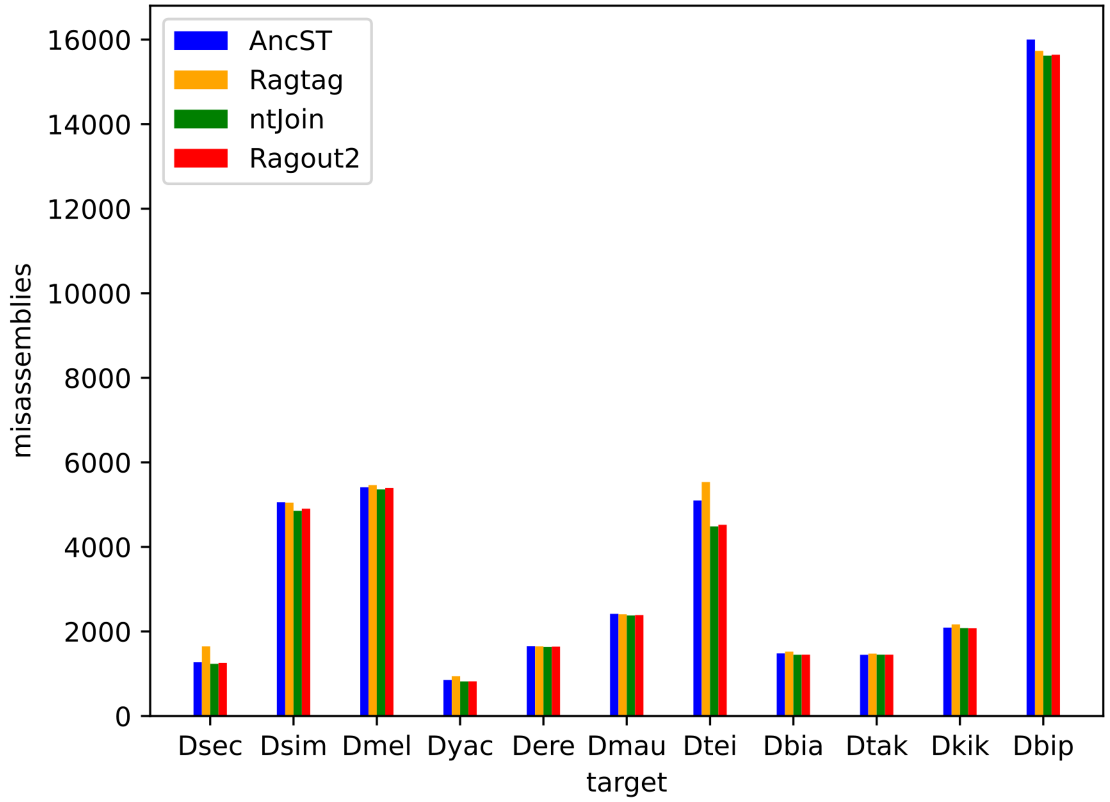


**Fig. 12** Number of *misassemblies* as computed by `Quast` for the 11 `Drosophila` newly scaffolded species with a chromosome- scale reference genome on NCBI. Further details in Table 3.

#### B.3 Assessment of Reference Chromosome Coverage with New Scaffolds

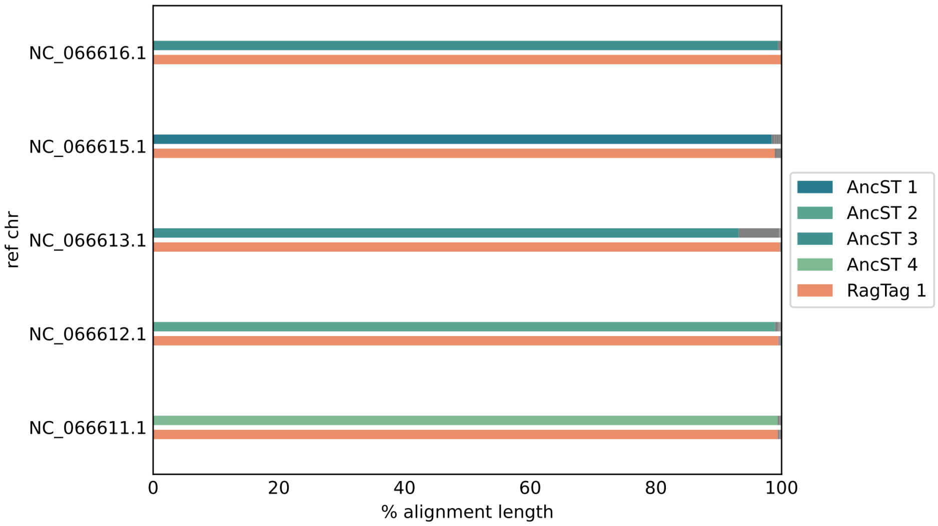


**Fig. 13** Shown are all reference chromosomes for the `Drosophila biarmipes` official reference assembly on NCBI on the y-axis. For each reference chromosome, the upper bar displays results computed with the `AncST`-based pipeline and the lower bar the ones from `RagTag`. The bars are stacked according to the coverage of each reference chromosome by new scaffolds from the respective tool. The coverage is estimated by the total alignment length of all `minimap` alignments recorded in the output of `Quast`. Only contigs/scaffolds covering at least a third of the total alignment length are drawn colored while the rest is kept gray. Each color represents a different new scaffold which are indicated in the legend.

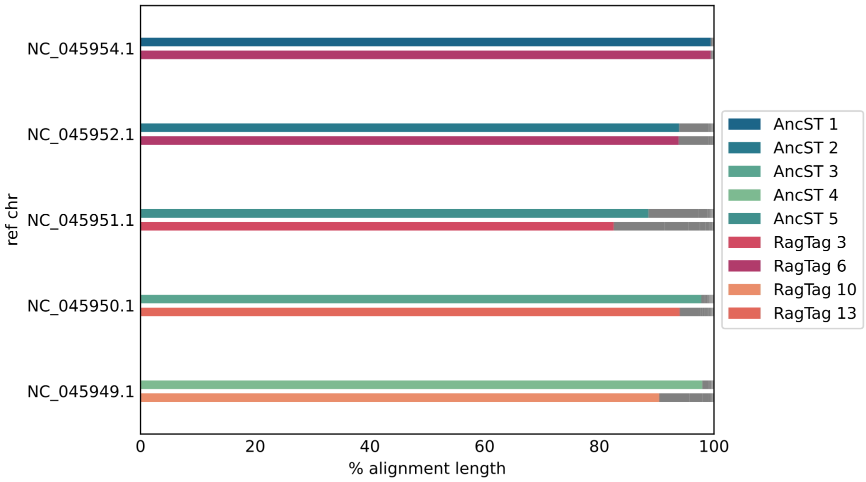


**Fig. 14** Assembly: GCA_018904445.1. Shown are all reference chromosomes for a species' official reference assembly on NCBI on the y-axis. For each reference chromosome, the upper bar displays results computed with the `AncST`-based pipeline and the lower bar the ones from `RagTag`. The bars are stacked according to the coverage of each reference chromosome by original contigs or new scaffolds from the respective tool. Each color represents a different original contig or new scaffold which are indicated in the legend. Only contigs/scaffolds covering at least a third of the total alignment length are drawn colored while the rest is kept gray. The coverage is estimated by the total alignment length of all `minimap` alignments recorded in the output of `Quast`. `From A.` means as computed by `AncST` and `From R.` means as computed by `RagTag`.

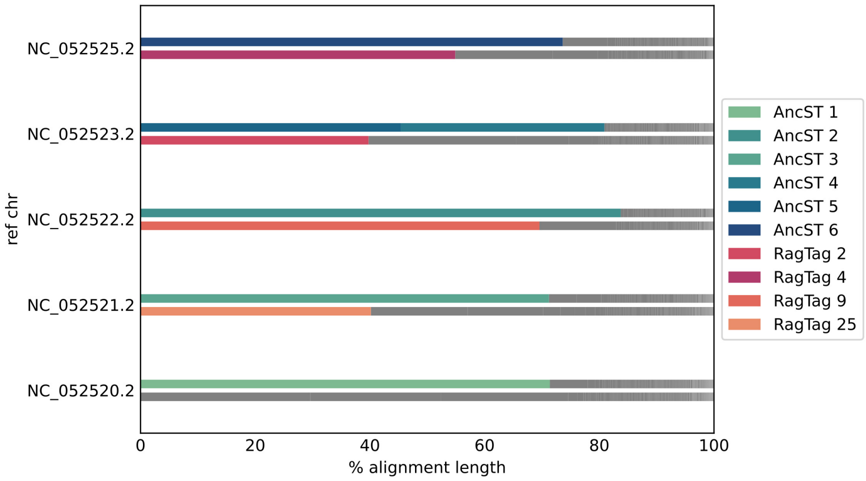


**Fig. 15** Assembly: GCA_039725655.1. See caption of Fig. 14.

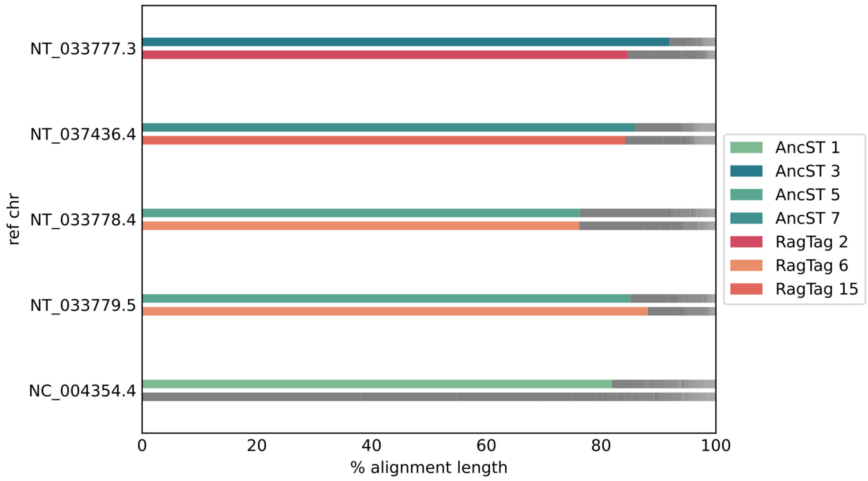


**Fig. 16** Assembly: GCA_000778455.1. See caption of Fig. 14.

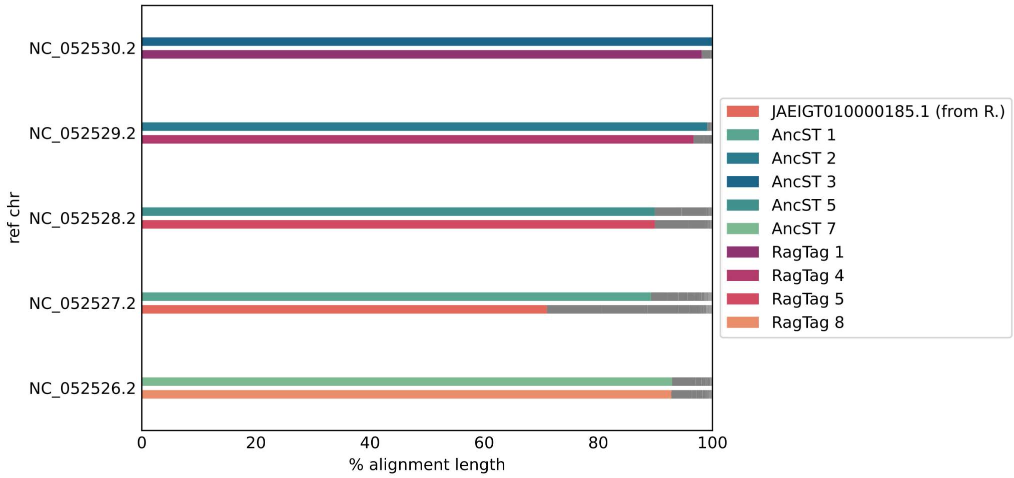


**Fig. 17** Assembly: GCA_018904385.1. See caption of Fig. 14.

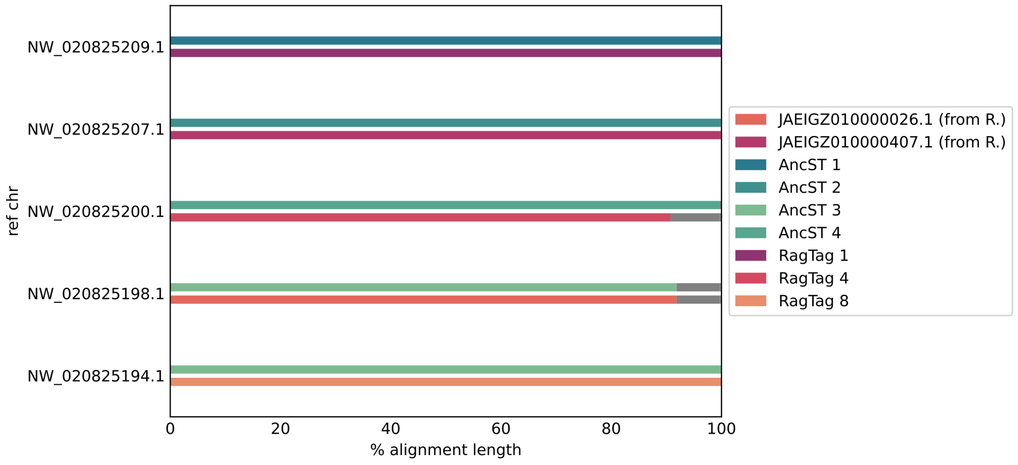


**Fig. 18** Assembly: GCA_018904525.1. See caption of Fig. 14.

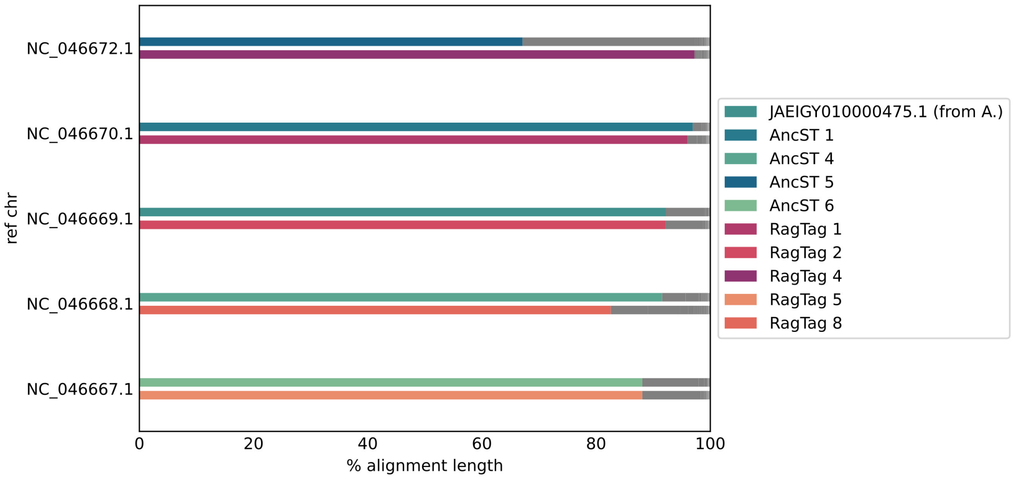


**Fig. 19** Assembly: GCA_018904475.1. See caption of Fig. 14.

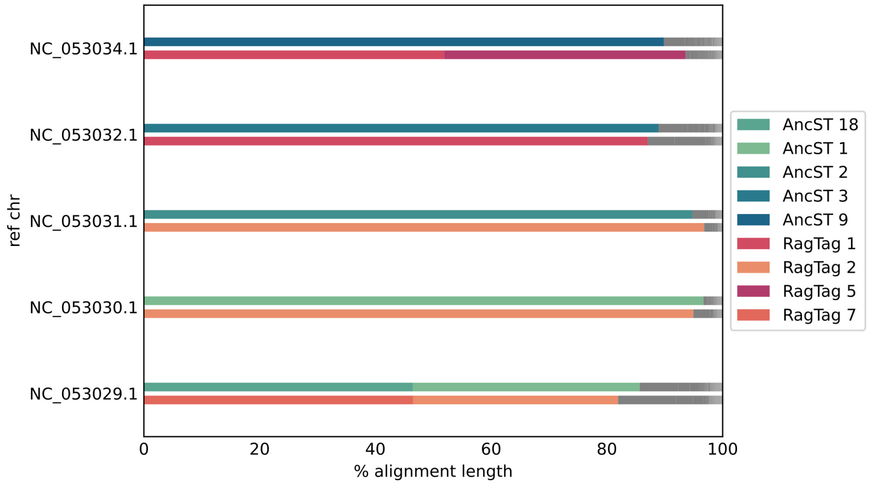


**Fig. 20** Assembly: GCA_018903625.1. See caption of Fig. 14.

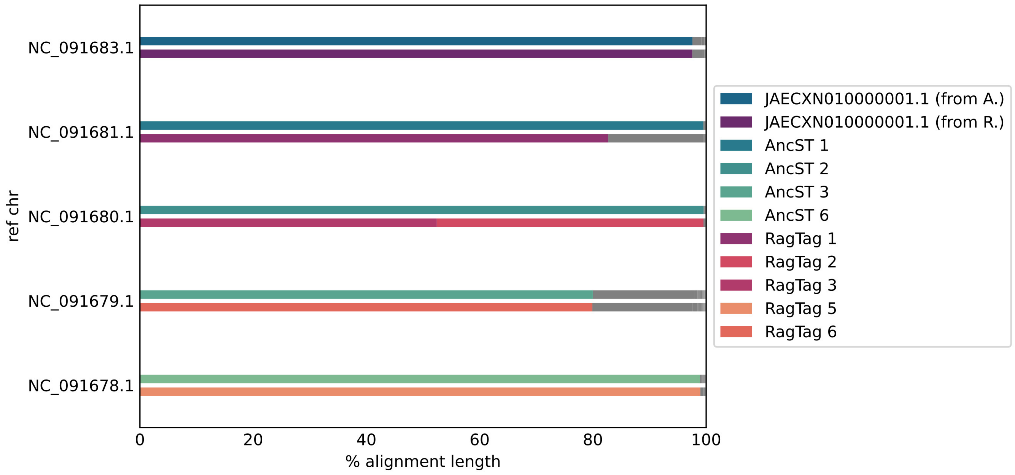


**Fig. 21** Assembly: GCA_018152695.1. See caption of Fig. 14.

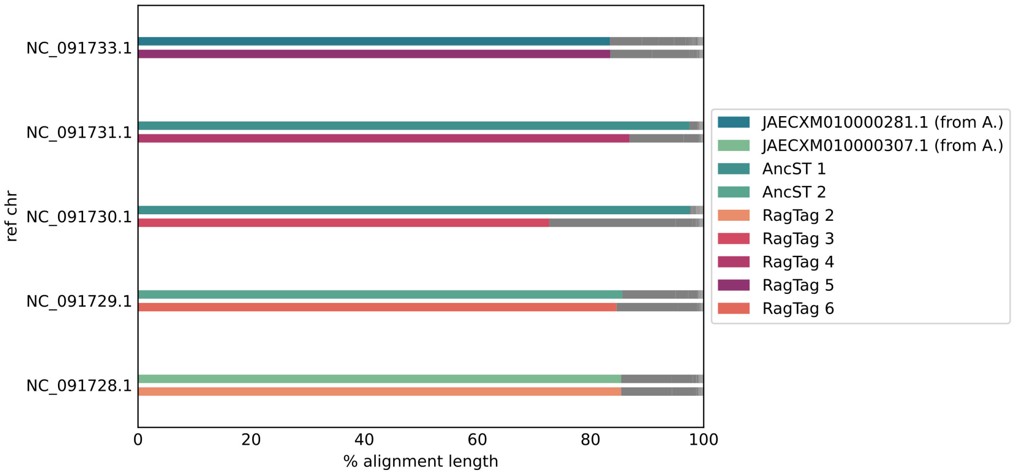


**Fig. 22** Assembly: GCA_018152535.1. See caption of Fig. 14.

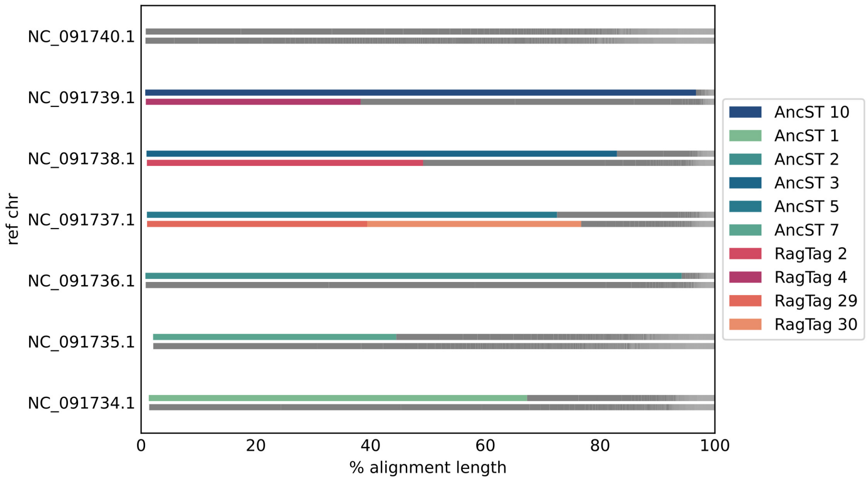


**Fig. 23** Assembly: GCA_021223765.1. See caption of Fig. 14.

#### B.4 Influence of Using Weights for RagTag and ntJoin on auN and Number of Misassemblies for Drosophila Scaffolds

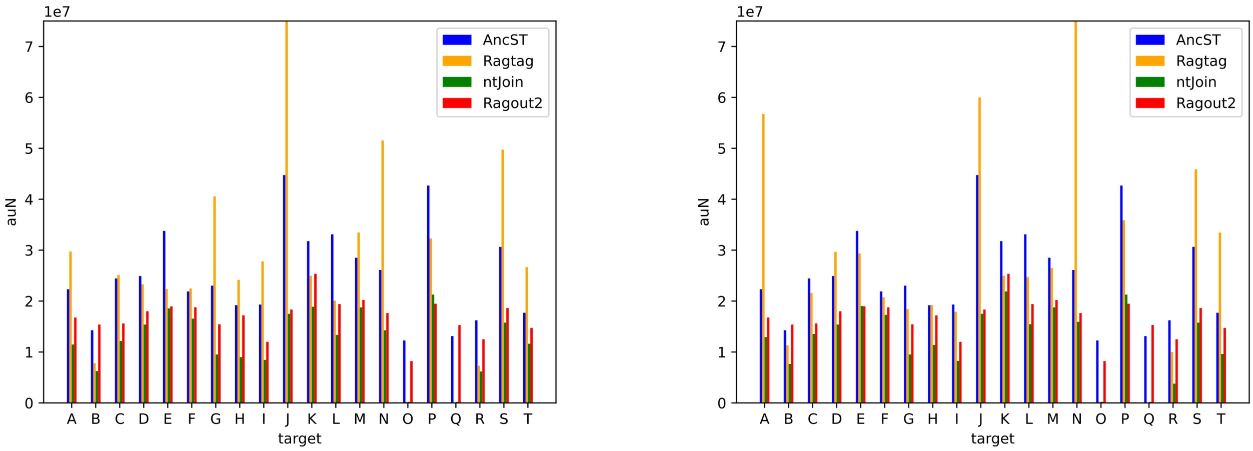


**Fig. 24** Left: Same plot as Fig. 11. Right: Same analysis but using `AncST` weights for `RagTag` and `ntJoin`.

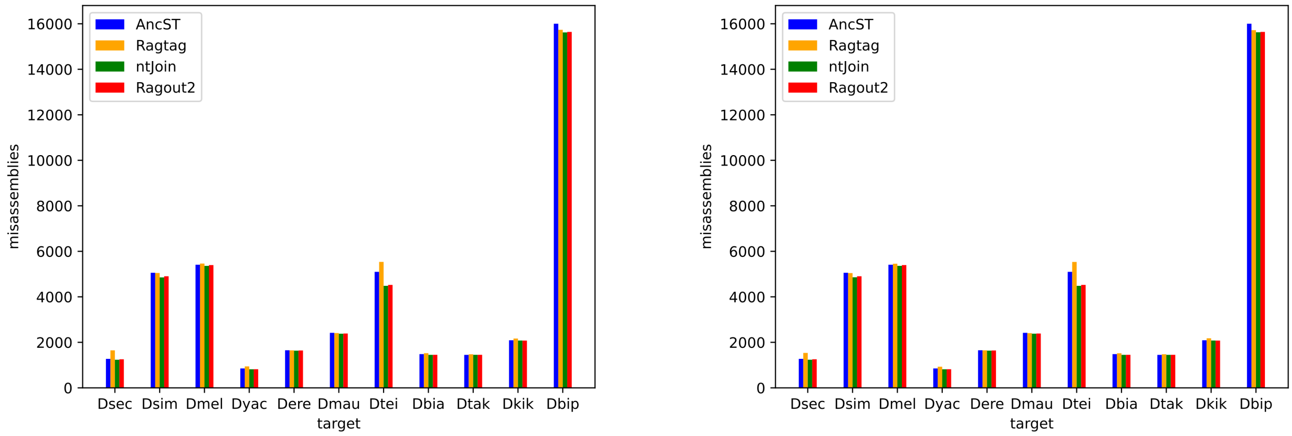


**Fig. 25** Left: Same plot as Fig. 12. Right: Same analysis but using `AncST` weights for `RagTag` and `ntJoin`.
(yes, the two plots are different)

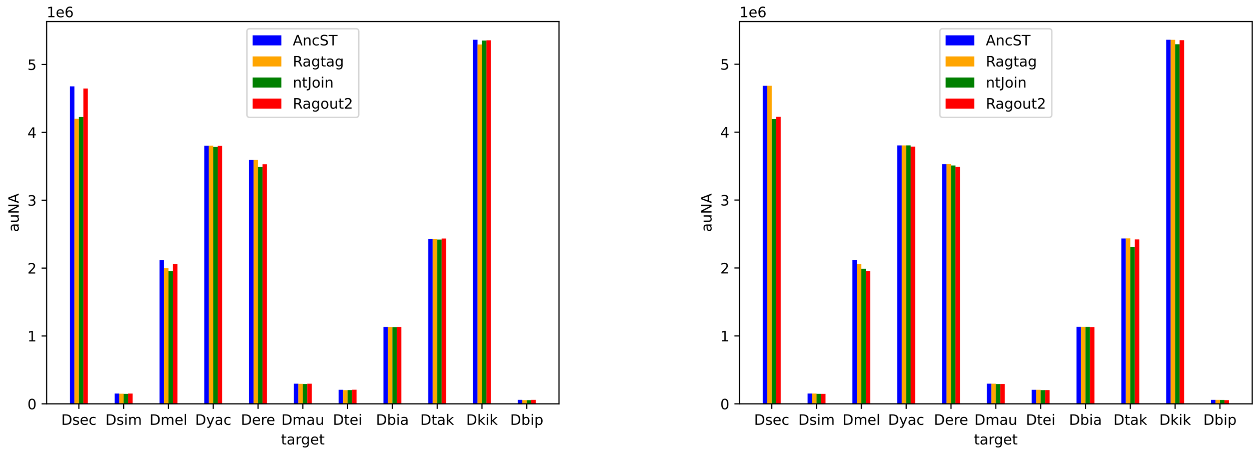


**Fig. 26** Left: auNA from `Quast` for each `Drosophila` species with a chromosome-level reference as in main text. Right: Same analysis as in main text but using `AncST` weights for `RagTag` and `ntJoin`.

### C Details of Tool Versions and Parameters

#### C.1 Installations of Scaffolders

`CSAR` and `Multi-CSAR` were installed from their sources[1]. Default parameters were used with the option `--nuc` to run on the nucleotide level on the two pairs of genomes as well as both reference genomes for the multi-reference version.
`RagTag` was installed from its source[2] and run on the two pairs of genomes. To obtain a multi-reference version we used its `merge` command on the result of the pairwise results. Default parameters were used. For the real test data `RagTag` was run with the `minimap` setting *asm 20* to increase sensitivity as we compare genomes of considerable divergence. `ntJoin 1.15` was installed using the respective `bioconda` recipe and was run with the following options to ensure comparability of the results: `agp=True no_cut=True time=True overlap=False`. `Ragout 2.3` was installed using the respective `bioconda` recipe and was run with the following options: `--refine --solid_scaffolds`. Multiple sequences alignments were generated with `cactus 2.9.9` and used in `maf` format as input for `Ragout2`. We used 58 cores for `AncST` anchor computation while the other tools were run as single-threaded applications. `CSAR` cannot be run multi-threaded although the consumed `user` time indicates use of multiple cores. We decided not to benchmark `RagTag`, `Ragout2` and `ntJoin` with multiple threads as they are so fast that time consumption is negligible already without multithreading. Multiple alignment computation with `cactus`, however, was done using 64 available cores.

[1] https://github.com/ablab-nthu/CSAR, https://github.com/ablab-nthu/Multi-CSAR
[2] https://github.com/malonge/RagTag

#### C.2 Details AncST Run

Both the primate and `Drosophila` genomes were processed at an instance of the `AncST` webserver available at https://anchored.bioinf.uni-leipzig.de with default parameters. That means that for k-mer statistics we calculate the k-0 mappability with $k = \frac{\log(genome\ size)}{\log(4)}$ with `GenMap`, we use initial candidate window sizes of 300 with a pitch of 50 and take the candidates up to the 42nd percentile of lowest aggregate k-mer counts. `macle` is used to calculate the genomes' match complexity and the highest scoring 42nd percentile of windows of size 1000 with pitch 50 is used. We did not use the feature to resolve duplicates of a copy low number as additional anchor candidates for this study (for details see webserver documentation and help).

#### C.3 Details Weights of Multi-ref AncST Scaffolding

As a weight for each single-reference scaffolding result we use the total `blast bit score` of all `AncST` alignments times 1000 divided by the length of the respective reference genome (to produce weights in a range of few dozens to few hundreds for human-readability). Accordingly, for each contig pair connected in one of the single-reference scaffolds, the produced weight is added to the corresponding edge in the superimposed graph for multi-reference scaffolding.

#### C.4 Robustness Single References

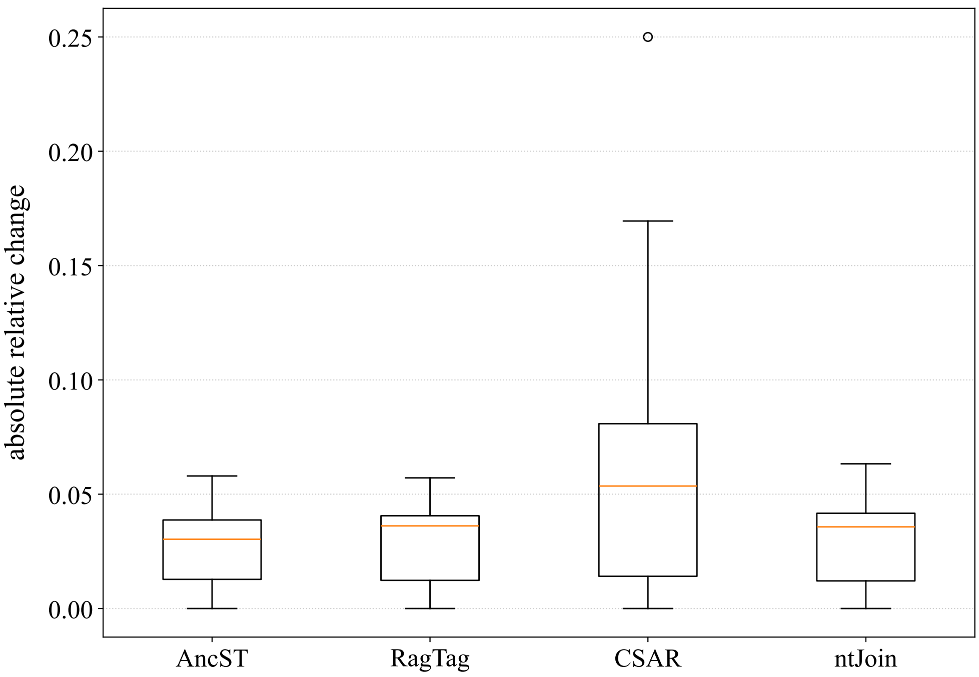


**Fig. 27** Shown are relative changes for all performance measures between using bonobo and chimp as references.

#### C.5 Implementation Details of Benchmarked Tools

**Table 4** Comparison of algorithms of scaffolding tools. Rows are tools; columns are the three stages (*Precomputation* for marker / alignment generation; *Single-ref scaffolding* for the single-reference contig ordering step when applicable, otherwise this column is also used for *Precomputation*; *Multi-ref scaffolding* for the reconciliation step over multiple references).

| Tool | Precomputation | Single-ref scaffolding | Multi-ref scaffolding |
|---|---|---|---|
| `AncST` | $d_0$-unique anchor candidates per genome; pairwise matches by $d_0 - \sqrt{\alpha\ell - d_0}$ distance cutoff ([2];this paper, Sec. Methods) | Greedy: contig assigned to its best-matching reference chromosome, then appended once $\geq 50\,\%$ of its anchors are visited along the reference; orientation by majority strand (this paper, Sec. Methods) | Max-weight perfect matching on the contig-end graph (`Blossom V` with 0-weight for *perfect* matching); cycles broken by min-weight edge removal (this paper, Sec. Methods) |
| `CSAR` / `Multi-CSAR` | Pairwise `NUCmer` or `PROmer` (`MUMmer`) alignment to obtain conserved genetic markers (see e.g. [3]); the `delta-filter -1` utility of `MUMmer` removes repeated markers from both target and reference (1-to-1 best mapping, [4], Sec. Implementation) | Algebraic-rearrangement framework: contig joining maximises the number of cycles in the marker adjacency graph (algebraic distance counted over reversals, transpositions and translocations); near-linear-time algorithm built on permutation groups and a disjoint-set data structure ([5], Sec. 2–3) | Max-weight perfect matching on the contig-end graph (`Blossom V` with 0-weight edges for *perfect* matching); cycles broken by min-weight edge removal ([6], Sec. Methods; [7]) |
| `RagTag` (incl. `merge`) | Pairwise `Minimap2`/`Unimap`/`Nucmer` alignment; unique-anchor filter as in [8], Sec. 2: default $\geq 10$ kbp of contig sequence not covered by any other alignment of that contig; consecutive same-strand alignments merged within 100 kbp ([9], Sec. RagTag whole-genome alignment filtering and merging) | Each contig assigned its primary alignment (longest merged alignment); contigs sorted by primary-alignment reference coordinate; oriented by primary alignment ([9], Sec. RagTag scaffold) | Max-weight matching on the scaffold graph (`NetworkX`, *not* perfect by default); cover graph augmented with $\infty$-weight intra-contig edges; min-weight edge removed from each cyclic connected component ([9], Sec. RagTag merge) |
| `ntJoin` | Ordered minimizer sketches per input; minimizers retained only if unique within each assembly *and* present in every assembly ([10], Sec. Materials and methods) | | Per branching with node degree $> 2$, incident edges are removed in increasing weight order until that node's degree drops to $\leq 2$ ([10], Sec. Materials and methods) |
| `Ragout 2` | `Cactus` MSA; iterative A-Bruijn graph at default scales 10 kb / 100 bp / 500 bp (largest scale forms the structural "skeleton"); bipartite max-weight matching of flanking-context blocks resolves repeat copies; alternating-cycle test removes chimeric/artificial edges; neighbour-joining phylogeny of the input genomes built from pairwise breakpoint distances ([11], Sec. Construction of synteny blocks; Sec. Synteny block size selection; Sec. Repeat resolution algorithm; Sec. Detection of chimeric adjacencies; Sec. Iterative assembly; Sec. Phylogenetic tree reconstruction) | | Synteny block ends are placed as vertices into a breakpoint graph and connected by colored edges with one reserved for known adjacencies from the target contigs. Each unknown contig-end vertex has potential candidates from synteny block adjacencies to any of the reference genomes and Sankoff DP over genome phylogeny is used to determine costs for each state. `Blossom` algorithm is applied to obtain the target adjacencies. ([7], [12], [11]) |
